# Molecular patterns of evolutionary changes throughout the whole nervous system of multiple nematode species

**DOI:** 10.1101/2024.11.23.624988

**Authors:** Itai Antoine Toker, Lidia Ripoll-Sánchez, Luke T. Geiger, Karan S. Saini, Isabel Beets, Petra E. Vértes, William R. Schafer, Eyal Ben-David, Oliver Hobert

## Abstract

One avenue to better understand brain evolution is to map molecular patterns of evolutionary changes in neuronal cell types across entire nervous systems of distantly related species. Generating whole-animal single-cell transcriptomes of three nematode species from the *Caenorhabditis* genus, we observed a remarkable stability of neuronal cell type identities over more than 45 million years of evolution. Conserved patterns of combinatorial expression of homeodomain transcription factors are among the best classifiers of homologous neuron classes. Unexpectedly, we discover an extensive divergence in neuronal signaling pathways. While identities of neurotransmitter-producing neurons (glutamate, acetylcholine, GABA and several monoamines) remain stable, ionotropic and metabotropic receptors for all these neurotransmitter systems show substantial divergence, resulting in more than half of all neuron classes changing their capacity to be receptive to specific neurotransmitters. Neuropeptidergic signaling is also remarkably divergent, both at the level of neuropeptide expression and receptor expression, yet the overall dense network topology of the wireless neuropeptidergic connectome remains stable. Novel neuronal signaling pathways are suggested by our discovery of small secreted proteins that show no obvious hallmarks of conventional neuropeptides, but show similar patterns of highly neuron-type-specific and highly evolvable expression profiles. In conclusion, by investigating the evolution of entire nervous systems at the resolution of single neuron classes, we uncover patterns that may reflect basic principles governing evolutionary novelty in neuronal circuits.

## Introduction

Evolutionary changes that shaped animal nervous systems played a pivotal role in their success in adapting to diverse environments. Characterizing these changes has been the goal of numerous comparative studies that enlightened our understanding of brain evolution. Classifications of neuronal cell types, a fundamental aim in neuroscience, traditionally relied mostly on anatomical features and, later, on electrophysiology and key molecular features such as neurotransmitter usage. More recently, these classifications have been further expanded and refined in various organisms through in-depth molecular characterization permitted by single-cell technologies (Tasic *et al*. 2016; Romanov *et al*. 2017; Sebe-Pedros *et al*. 2018; Zeisel *et al*. 2018; Allen *et al*. 2020; Deryckere *et al*. 2023). When applied comparatively in different species, these techniques can reveal evolutionary novel neuronal types which arose in a particular species or taxon, the species-specific loss of a neuron-type or the rare presence of a homologous cell type previously thought to be lacking in a species (Krienen *et al*. 2020; Yamagata *et al*. 2021; Schmitz *et al*. 2022; Shafer *et al*. 2022; Wei *et al*. 2022; Chen *et al*. 2023; Hahn *et al*. 2023; Lamanna *et al*. 2023; Ye *et al*. 2024). Molecular profiling also traces the evolutionary trajectories of neuronal cell classes or brain regions across divergent species, revealing whether neuronal features with functional similarities resulted from homology or from convergent evolution (Horie *et al*. 2018; Tosches *et al*. 2018; Colquitt *et al*. 2021; Hain *et al*. 2022; Woych *et al*. 2022).

In-depth analyses of homologous neuron types across species enables the characterization of diverging and conserved molecular features with the potential to address critical open questions about brain evolution (Roberts *et al*. 2022). For example, do the patterns of deployment of different modes of neuronal communication (different neurotransmitter systems, synaptic vs extrasynaptic etc.) diverge in their evolutionary trajectories? Is evolutionary change reflected uniformly in the entire brain, or conversely do specific regions, circuits or neuron features constitute evolutionary hotspots? Shining light on these central questions in vertebrate brains presents several challenges. The sheer size of brains often necessitates the characterization of specific regions in isolation, preventing a global investigation across the entire nervous system with minimal sampling bias. Moreover, cataloguing the complete collection of neuronal classes (both molecularly and anatomically) and the precise sets of regulatory factors specifying their terminal identity, a key requirement when defining homologous neurons (Arendt *et al*. 2016; Arendt *et al*. 2019), is still an ongoing endeavor in complex organisms.

While the complexity of vertebrate brains complicates the analysis of evolutionary trajectories on a whole nervous system level, the compact nervous system of nematodes (Schafer 2016) permits the identification of nervous system-wide patterns and principles of evolutionary change through comparative analysis. Overall anatomical organization and embryonic lineaging studies suggest that the nervous systems of distinct nematodes may be very similar (Zhao *et al*. 2008; Schafer 2016; Memar *et al*. 2019). However, this notion has never been rigorously tested with proper cellular and molecular resolution. We interrogate this issue here using three *Caenorhabditis* species, *C. elegans, C. briggsae* and *C. tropicalis,* that are thought to have shared their last common ancestor more than 45 million years ago (Figure 1A)(Qing *et al*. 2023)(for comparison, humans and chimpanzees diverged less than 10 million years ago). Nematodes from the *Caenorhabditis* genus exhibit interspecies biases towards different climates and ecological niches, and display a plethora of differences in behavioral patterns (Chasnov *et al*. 2007; Kiontke *et al*. 2011; Felix and Duveau 2012; Felix *et al*. 2013; Stegeman *et al*. 2013; Poullet *et al*. 2015; Burton *et al*. 2021; Crombie *et al*. 2022; Crombie *et al*. 2024; Ebert and Bargmann 2024).

**Figure 1.**
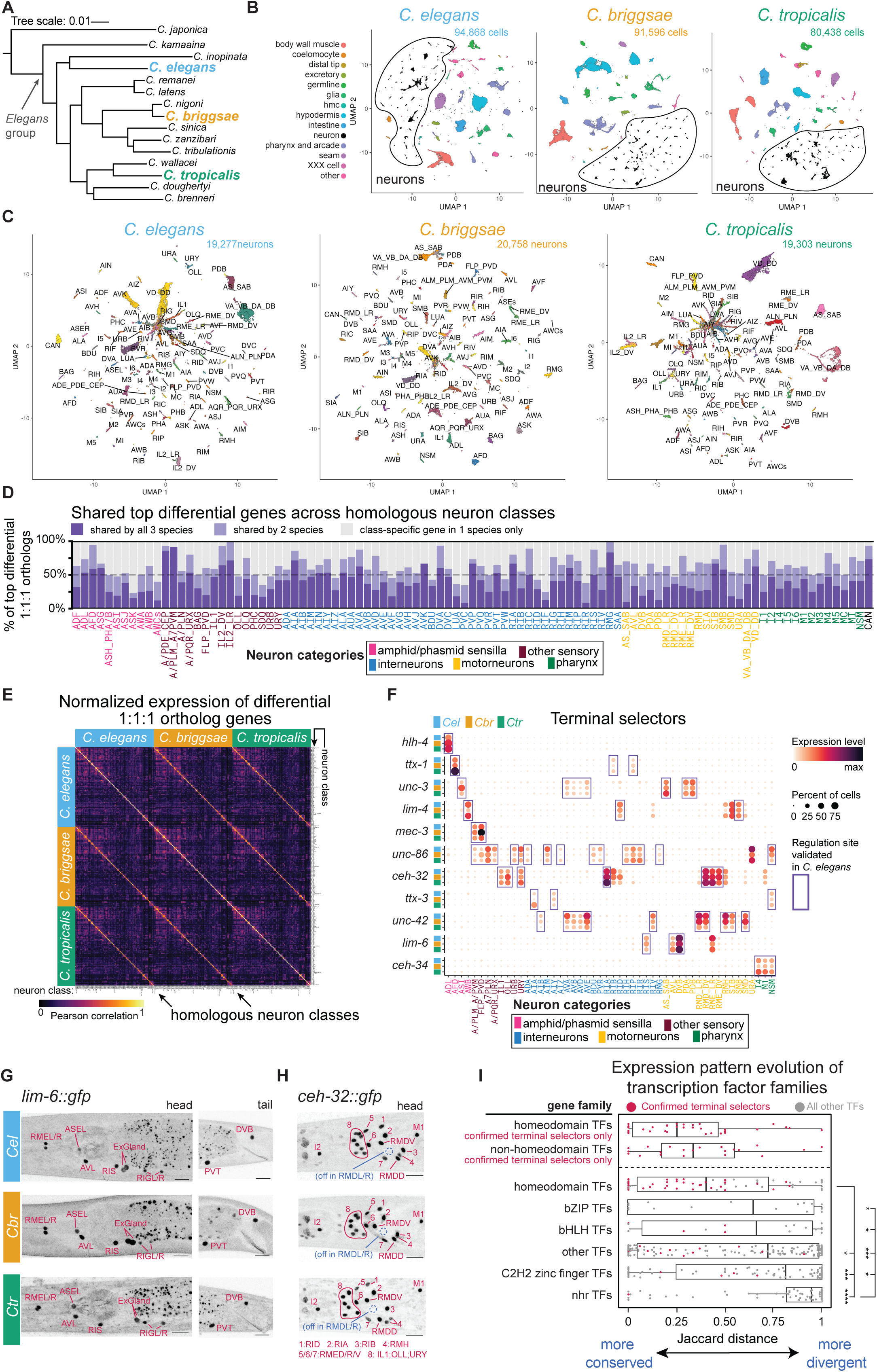
scRNA atlases of 3 nematode species delineate homologous neuronal cell types with conserved expression of neuronal identity specifiers. (A) Phylogenetic tree of species in the *Elegans* group, adapted from (Stevens *et al*. 2020). Scale: substitutions per site. (B) UMAP representations of sequenced cells colored by tissue in 3 *Caenorhabditis* species. (C) UMAPs of all neuronal cells. Neuron identities were determined based on 1:1:1 ortholog genes differentially-expressed in *C. elegans*. (D) Barplots representing all 1:1:1 orthologs (y-axis) in the top-10 of differential genes of a neuron class in any of the three species. Colors within bars depict the gene subsets found in the top-20 differential genes in all three species (dark purple), in two species (light purple) or in one species (grey). (E) Correlation matrix of pseudobulked normalized expression levels of 1,380 differentially-expressed 1:1:1 orthologs (marker_score >0.1 in at least one species and one neuron class). Rows and columns are grouped by species, and then by neuron class. (F) Cross-species expression dotplot for a subset of transcription factor orthologs functionally validated as identity specifiers in *C. elegans*. Full list of genes and neurons are shown in Figure S2. Nematode species (y-axis) and neuron class (x-axis) are color-coded according to legend. Dot size represents the fraction of cells expressing the gene in a given neuron class, color represents scaled average expression levels. **(G-H)** Representative fluorescent microscopy images (Z-stack max projections) of *lim-6::gfp.* (G) and *ceh-32::gfp* (H) protein fusions in nematodes genomically tagged in the endogenous loci of the corresponding transcription factor. Names of expressing neuron classes are indicated in red. Scale bars: 10μm. **(I)** Jaccard distances of transcription factors from different families. Each dot represents a gene. Red dots represent genes that are experimentally validated specifiers of terminal identity of at least one *C. elegans* neuron class. Boxplots are Tukey-style. Top two rows display separately the terminal selector genes alone (these genes thus appear twice in the plot) for ease of visualization and are not included in statistical tests. Kruskal-Wallis test with Dunn’s correction. \*\*\*\**P* < 0.0001, \*\*\**P* < 0.001, \**P* < 0.025, unmarked comparisons were not significant.

We hypothesized that using the knowledge accumulated from *C. elegans* neuronal development, anatomy and function, we would be able to assess the extent to which cell types can be homologized across the *Caenorhabditis* genus and to characterize the evolution of gene expression signatures of an entire nervous system at the resolution of single neuronal classes. On the basis of our single cell transcriptomic data, validated by genome-engineered fluorescent reporter alleles in all three nematode species (25 generated for this particular study), we provide an in-depth analysis of different gene families involved in neuronal fate specification and neuronal signaling. We discovered striking patterns of evolutionary stability as well as evolutionary change in homologous neuronal cell types. We assessed whether the changes are enriched in subsets of neurons or homogenously distributed in the nervous system. Finally, we present a primary characterization of a family of small, secreted proteins of unknown function, that are abundantly expressed in the nervous system in a neuron class-specific manner. Taken together, our analysis reveals striking patterns of evolutionary changes in neuronal signaling across 45 millions years of evolution, distributed throughout the entire nervous system of each nematode species.

## Results

### scRNA atlases of three *Caenorhabditis* species delineate homologous neuronal cell types with conserved expression of neuronal identity specifiers

We generated scRNA-seq libraries from the three androdiecious *Caenorhabditis* species *C. elegans, C. briggsae* and *C. tropicalis* at the second larval stage (L2) of development (Figure 1A). At this stage, all neuronal cells have been generated and wired up into a fully functional nervous system (Sulston 1983; Witvliet *et al*. 2021). After quality filtering, we obtained 94,868 *C. elegans* cells, 91,596 *C. briggsae* cells and 80,438 *C. tropicalis* cells for a total of 266,902 sequenced and tissue-annotated single cells (Figure 1B and table S1). Cell clusters were assigned to organ tissues based on examination of cluster-enriched differentially-expressed genes with 1:1:1 primary sequence orthologs across the three species and comparisons with prior tissue-specific transcriptome studies in *C. elegans* (see **Methods**)(Cao *et al*. 2017; Packer *et al*. 2019; Taylor *et al*. 2021).

We subclustered neuronal cells and refined the annotation of neurons into molecularly distinct neuronal classes in the three species (Figure 1C). The final neuronal datasets include 59,338 neurons (ranging 19,277∼20,758 per species) encompassing 114/118 (*C. elegans* & *C. briggsae*) and 112/118 (*C. tropicalis*) neuron classes. “Missing” neuronal cell classes (HSN, RMF, PVN, VC, I3, ASE) do not stem from losses of the neuron types in either nematode species since all these classes were reliably detected using CRISPR/Cas9-engineered reporter alleles for neurotransmitter pathway genes (as described further below). Detailed information about sequenced cells from all neuronal groups and species appear in table S1.

We were able to robustly assign homology to all sequenced neuron classes based on the following arguments: First, large subsets of the most differentially expressed 1:1:1 ortholog genes were shared across species in most cell types (Figure 1D, and table S2): in 83% of all neuronal cell groups, at least 50% of the top-10 cell-specific differential genes in any species were present among the top-20 differential genes in another species. Even in the extreme lowest case (ASK sensory neurons), this proportion was still of 23% top-10 differential genes shared by at least two species (Figure 1D). Second, hierarchical clustering of cell types by the expression of variable genes showed high concordance with our marker-based cell-type calls (Figure 1E and S1). Third, when considering cross-species transcriptomic data, an accepted minimal definition for homologous cell types is that they will share a conserved set of transcription factors specifying their terminal identity (Arendt *et al*. 2019). We expanded this concept further by specifically focusing on transcription factors that are proven key regulators of neuronal identity in *C. elegans* (including so-called “terminal selectors”). Such critical regulatory factors were determined for 111 out of the 118 neuronal classes through extensive genetic mutant analysis (table S3)(Hobert 2016; Reilly *et al*. 2022). To date, the *C. elegans* neuronal “regulatory code” includes 62 transcription factor-encoding genes acting as identity specifiers in at least one neuron class, 60 of which have 1:1:1 orthologs in our gene models. Cross-species expression data for this set of transcription factors displays overwhelmingly conserved cell-specific expression across species in the homologous cell types for which they act as terminal selectors in *C. elegans* (Figure 1F and S2).

We experimentally validated the conservation of neuronal identity-specific transcription factors by using CRISPR/Cas9 to insert *gfp* in the genomic loci of two such terminal selectors genes, the LIM homeodomain gene *lim-6* and the SIX3/6-like homeodomain gene *ceh-32.* We had previously described the expression patterns of these genes in *C. elegans* (Reilly *et al*. 2020) and by now tagging their orthologs in *C. briggsae* and in *C. tropicalis,* we confirmed conserved neuron-class-specific expression using fluorescent microscopy (Figure 1G-H). These results strengthen the neuron class homology assignments by grounding them in regulatory programs of neuronal differentiation.

Neuronal cells that share extensive phenotypic similarity and belong to the same neuron type can sometimes be further subclassified into neuronal subtypes when they differ in only a small number of reproducible characteristics (White *et al*. 1986; Hobert *et al*. 2016). For example, the RMD-class neck motor neurons include three bilateral neuron pairs (dorsal, lateral, and ventral) that share numerous anatomical and molecular features in common, yet the dorsoventral pairs differ from the lateral pair in the expression of several terminal features such as the glutamate AMPA receptor *glr-2* and the neuropeptides *nlp-11* and *flp-19* (Cros and Hobert 2022). Our analysis of the scRNA-seq datasets captured molecularly distinct subclusters of RMD-, IL2- and RME-class neurons across all *Caenorhabditis* species (Figure S3). In *C. elegans* RMD, subclass-specific effector gene expression is regulated by *ceh-32* which is expressed in dorsoventral but not lateral RMD neurons, and act as a specifier of RMD dorsoventral subtype identity (Cros and Hobert 2022). Similarly, *lim-6* is expressed in lateral but not dorsoventral RME motoneurons, and in left but not right ASE sensory neurons (Hobert *et al*. 1999). Using our endogenous protein fusion reporters, we found that the subtype-specific expression patterns of CEH-32 and LIM-6 are conserved also in *C. briggsae* and *C. tropicalis* (Figure 1G-H). Together, these findings underscore the extent by which cell type classification is conserved across many million years of nematode phylogeny.

### Homeodomain transcription factor codes as classifiers of neuronal cell types

After establishing the class identity of neuronal clusters, we characterized the evolutionary dynamics of gene families that define the functional properties of neuron classes. First, for every gene and every neuron class, we sought to determine whether the detected signal from sequencing reads reflects high-confidence true expression. To achieve this, we trained a random forest classifier on our *C. elegans* dataset and 14,406 neuron-type-specific “ground truth” observations from 147 genes that were previously validated with fluorescent markers (CRISPR-tagged endogenous genes or fosmid-based reporters, table S4). We then applied the newly-generated classifier on all three datasets and obtained thresholded data of binary (ON/OFF) expression values. An analogous thresholding approach (although not machine learning-based) was successful in past *C. elegans* scRNA-seq studies and was even able to unveil true sites of expression that had been previously missed in transgenic reporter constructs limited in their cis-regulatory information (Cao *et al*. 2017; Packer *et al*. 2019; Taylor *et al*. 2021; Wang *et al*. 2024). We integrated the thresholded expression data across species and neuron types to calculate the Jaccard distances for every 1:1:1 orthologous gene triplet, reflecting how evolutionary divergent the gene is in its neuron-class-specific expression across species (table S4). Genes displaying similar (conserved) neuron-type-specific expression patterns in all three species have lower Jaccard distance values than (divergent) genes expressed in different sets of neurons in different species.

We found that the Jaccard distances of homeobox genes were more conserved than any other family of transcription factors (Figure 1I). This result is consistent with the high prevalence of homeobox genes among confirmed terminal selectors of neuronal identity (Figure S2, table S3). When inquiring specifically the subset of 22 non-homeodomain transcription factors that are validated terminal selectors, the Jaccard distances for this group were as low (conserved) as the homeodomain family. At the other extreme of the conservation spectrum, the TF family of C4 zinc finger, nuclear hormone receptors stand out as particularly divergent in their neuron-type-specific expression across species. Compared to other TF families, members of the homeodomain TFs are the most sparsely expressed (i.e., in fewer neuron classes), yet their sparsity is sufficient to combinatorially define each individual neuron class (Figure S4). Taken together, thanks to their conservation and their relative sparsity, combinatorial homeobox gene expression profiles may represent the most robust classifiers of cell type identity across metazoans (Hain *et al*. 2022; Najle *et al*. 2023; Fung *et al*. 2024).

### Expression of species-specific genes and differential expression of conserved orthologs are two independent modes of evolutionary novelty differentially harnessed by sensory, motor and interneurons

Are certain neuron types more evolutionary labile than others in their usage of the pool of genes present in the genome? To answer this question, we adopted a cell-centered approach and dissected two different modalities of evolutionary change in neuron-type-specific gene expression. The first potential modality of divergence within homologous neurons is through the gain or loss of species-specific genes, and the second is through the differential expression of homologous genes.

Initially, we established sets of species-specific genes for each nematode, defined as genes with no sequence homology to genes in neither of the two other species (i.e. genes with “1-to-none” orthology relationships). We note that limiting the analysis to “1-to-none” genes is too restrictive to capture all species-specific genes, since the latter also include recent duplications of conserved genes, which we discuss further below. We found that the number of expressed “1-to-none” species-specific genes was consistently higher (21%∼92% increase) in ciliated neurons belonging to the amphid and phasmid sensory sensilla compared to any other neuron category in all species, both in absolute numbers and when normalized to the total number of expressed genes per neuron class (Figure 2A and S5). The amphid and phasmid sensilla constitute the main olfactory, chemosensory and thermosensory organs in *C. elegans*, and their neurons expressed more species-specific genes than sensory neurons sensitive to other modalities such as CO_2_, O_2_ or mechanosensation. Some additional statistically significant differences could be detected in comparisons between other functional categories of neuronal classes, but those differences were not consistent across all species and were always much smaller in their effect size (Figure 2A and S5).

**Figure 2.**
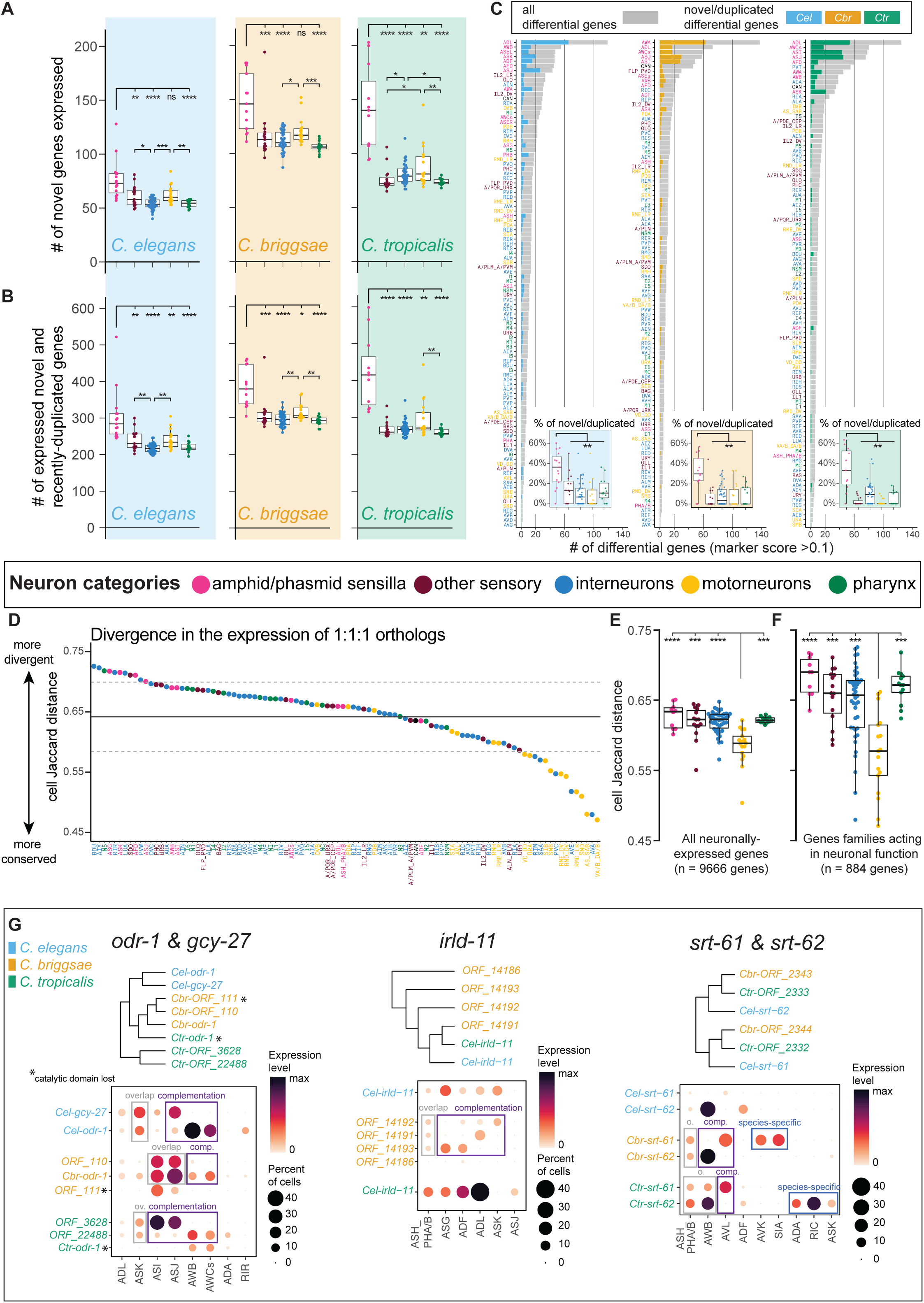
Distinct neuron functional categories differ in their preferred mode of transcriptional novelty. **(A)** Number of species-specific novel genes (“1-to-none” orthologs) expressed in different neuron classes and species. Neuron classes were grouped into functional categories (x-axis and color coded), each dot represents a single neuron class. Boxplots are Tukey-style. Kruskal-Wallis test with Dunn’s correction. **(B)** Number of species-specific novel genes and recently duplicated genes (“1-to-none”, “1-to-many”, “many-to-many” orthologs) expressed in different neuron classes and species. **(C)** Number of differentially-expressed genes (top_markers score >0.1, x-axis) expressed in each neuron class (y-axis) and species. Genes with high marker scores tend to be abundantly and specifically expressed in one or few neuron classes. Total numbers appear in grey bars, the subsets of species-specific novel genes and recently duplicated genes are colored according to the corresponding species. Bottom panels – Proportions of novel and duplicated genes out of all differentially-expressed gene per neuron class, grouped by neuron functional category. Each dot represent a neuron class. **(D)** Jaccard distances (y-axis) measuring the divergence in the usage of 1:1:1 ortholog triplets in each homologous neuron class (x-axis) across species. Neuron classes are ordered in decreasing Jaccard distance and color-coded according to functional categories. Horizontal bars: mean ± 1sd. **(E-F)** Jaccard distances (y-axis) of neuron classes (dots) grouped into functional categories (x-axis and colors). The calculation of Jaccard distances included (E) 8256 genes expressed in at least two species anywhere in the nervous system or (F) 785 genes belonging to gene families with established function in the nervous system (Hobert 2013). Kruskal-Wallis test with Dunn’s correction. \*\*\*\**P* < 0.0001, \*\*\**P* < 0.001, \*\**P* < 0.01, \**P* < 0.025. (G) Expression patterns for orthogroups composed of closely-related homologous genes. Genes are color-coded according to their species. Cases of paralog overlap, complementation and neofunctionalization are labeled in rectangles. Schematic gene phylogeny trees are shown above panels.

To account also for recently duplicated species-specific genes, we generated separate sets of species-specific genes which included “1-to-none” genes together with “many-to-1” and “many-to-many” gene mappings. These new “permissive” sets are only an approximation of the true pool of species-specific genes, since they include duplicated paralogs as well as the corresponding ancestral orthologs in all cases where the two could not be distinguished due to high sequence homology and high synteny. Remarkably, the “1-to-none” gene sets and the permissive gene sets displayed the very same evolutionary patterns of neuronal gene expression (Figure 2B and S5). We therefore deduce that these results are in high likelihood the patterns displayed by the bona fide sets of novel species-specific genes.

One family standing out among the sets of species-specific genes is the family of G-protein coupled receptor (GPCR)-encoding genes. GPCR genes are abundant and diverge rapidly in the genomes of *Caenorhabditis* nematodes and of many other animal clades, including mammals (Nei *et al*. 2008; Thomas and Robertson 2008; Ma *et al*. 2021; Ma *et al*. 2024). GPCRs clearly contributed to the enrichment of novel genes in olfactory and chemosensory neurons, since the vast majority of GPCRs are expressed in these neurons and since GPCRs constitute up to 19% of all novel genes expressed in the nervous system (Ma *et al*. 2024). Yet, the enrichment was still evident even when GPCRs were completely excluded from the expression analyses (Figure S5), indicating that GPCRs are not the sole contributors to the observed pattern. Alongside GPCRs, other species-specific genes enriched in amphid sensilla neurons included nuclear hormone receptors (nhr), Insulin/EGF Receptor-L Domain genes (irld), and genes with no identifiable protein domains, the latter being by far the most abundant gene category among neuronally-expressed species-specific genes (Figure S5).

Finally, species-specific genes were prominent among the top differentially-expressed genes in amphid and phasmid sensilla neurons (Figure 2C), even when excluding GPCRs (Figure S5). Hence, they are not merely expressed at detectable levels, but also tend to be abundantly and specifically expressed in those neuron classes. These findings suggest that the usage of species-specific genes is a frequent form of evolutionary novelty and a major feature of the molecular signature of chemosensory and olfactory neurons, while being less prominent in other categories of neurons.

Besides divergence in the usage of novel genes, homologous neuron types in evolving species can acquire species-specific molecular signatures through divergence in the expression of conserved, orthologous genes. To characterize this type of divergence, we calculated a Jaccard distance for each neuron class reflecting how divergent or conserved is the expression pattern of neuronally-expressed orthologous genes therein (Figure 2D). The pattern emerging from the analysis of orthologous genes differed sharply from the pattern of novel species-specific genes described above. We found that the expression of 1:1:1 orthologs was significantly more conserved in motor neurons compared to other neuron functional categories (Figures 2D-F). Importantly, sensory neurons were not particularly divergent through this metric, displaying divergence values similar to the values of interneurons or pharyngeal neurons. This was true when we included in the analysis either all neuronally-expressed 1:1:1 orthologs (8254 genes, Figure 2E) or a restricted pool of 785 1:1:1 orthologs belonging to gene families known to impact the functional properties of neurons (Figure 2F)(Hobert 2013).

In summary, our findings suggest that neuron groups are differentially impacted by two “arms” of gene expression divergences exhibiting separate evolutionary dynamics: ciliated olfactory/chemosensory neurons of the amphid and phasmid sensilla make extensive use of novel and rapidly duplicating genes, resulting in a more species-specific molecular signature compared to other categories of neurons. In parallel and in stark contrast, neuron-class-specific expression of 1:1:1 orthologs is more evolutionary conserved in motor neurons while the levels of divergence are comparable in all remaining neuron categories, including ciliated sensory neurons. Rates of evolutionary novelty in the neuronal usage of conserved genes show no obvious “hotspot” and are slower only in motor neurons.

### Subfunctionalization, neofunctionalization, overlap and degeneration of duplicated genes at single neuron resolution

Gene duplication is thought to play a major role in cellular innovation (Ohno 1970; Force *et al*. 1999; Nei *et al*. 2008). After duplication, redundant genes often undergo negative selection that result in degeneration and pseudogenization. In other cases, gene duplicates can diverge to adopt new functions (neofunctionalization) or to “divide labor” (subfunctionalization) by subsetting the ancestral gene function between two genetic loci now evolving independently. From a cell type perspective, neofunctionalization and subfunctionalization can arise in a species-specific manner when gene duplicates are differentially employed by homologous cell types. Exploration of our datasets enables the investigation of the expression patterns of paralogous expansions in the nervous system across species, highlighting several intriguing cases.

One example is the orthogroup that includes the guanylyl cyclase genes *odr-1* and *gcy-27* in *C. elegans* and 3 paralogs in both *C. briggsae* and *C. tropicalis*. In *C. elegans*, both *odr-1* and *gcy-27* take part in olfactory and gustatory response signaling pathways (Bargmann *et al*. 1993; Krzyzanowski *et al*. 2013). Considering all paralogs in each species together, this orthogroup is consistently expressed in the sensory neurons ASI, ASJ, ASK, AWB, AWCs, and to a lesser extent ADL, however different species display different patterns of paralog overlap and complementation (Figure 2G). In both *C. briggsae* and *C. tropicalis*, that have 3 paralogs, one of those paralogs lost its guanylate cyclase catalytic domain and is expressed at low mRNA levels in cells overlapping with another paralog, thus exhibiting a probable case of duplicate degeneration both at the level of enzymatic function and transcript expression. Moreover, all *C. briggsae* paralogs overlap their expression in ASI and ASJ neurons, while *C. elegans* and *C. tropicalis* display a subfunctionalization pattern in which ASI and ASJ are heavily biased towards one paralog, while the AWB and AWC neurons rely on the second paralog. Finally, species- and class-specific novel expression of genes in this *odr-1/gcy-27* orthogroup (i.e. neofunctionalization) was detected in *C. elegans* RIR and in *C. tropicalis* ADA interneurons. Analysis of two additional orthogroups, *irld-11* (Insulin/EGF receptor L domain protein) and *srt-61/62* (GPCRs), are depicted in Figure 2G.

### Evolutionary plasticity of neurotransmitter signaling pathways

Information flow in the nervous system is mediated by various chemical signals such as neurotransmitters, monoamines and neuropeptides. What are the evolutionary dynamics of the gene expression patterns of these neuronal signaling gene modules? Strikingly, we found that the expression patterns of genes involved in synthesis and vesicular loading of specific neurotransmitters (acetylcholine, GABA, glutamate) and monoamines (dopamine, serotonin, octopamine and tyramine) were very conserved, with conservation scores similar to cell-fate specifying transcription factors (Figure 3A-B and Figure S6).

**Figure 3.**
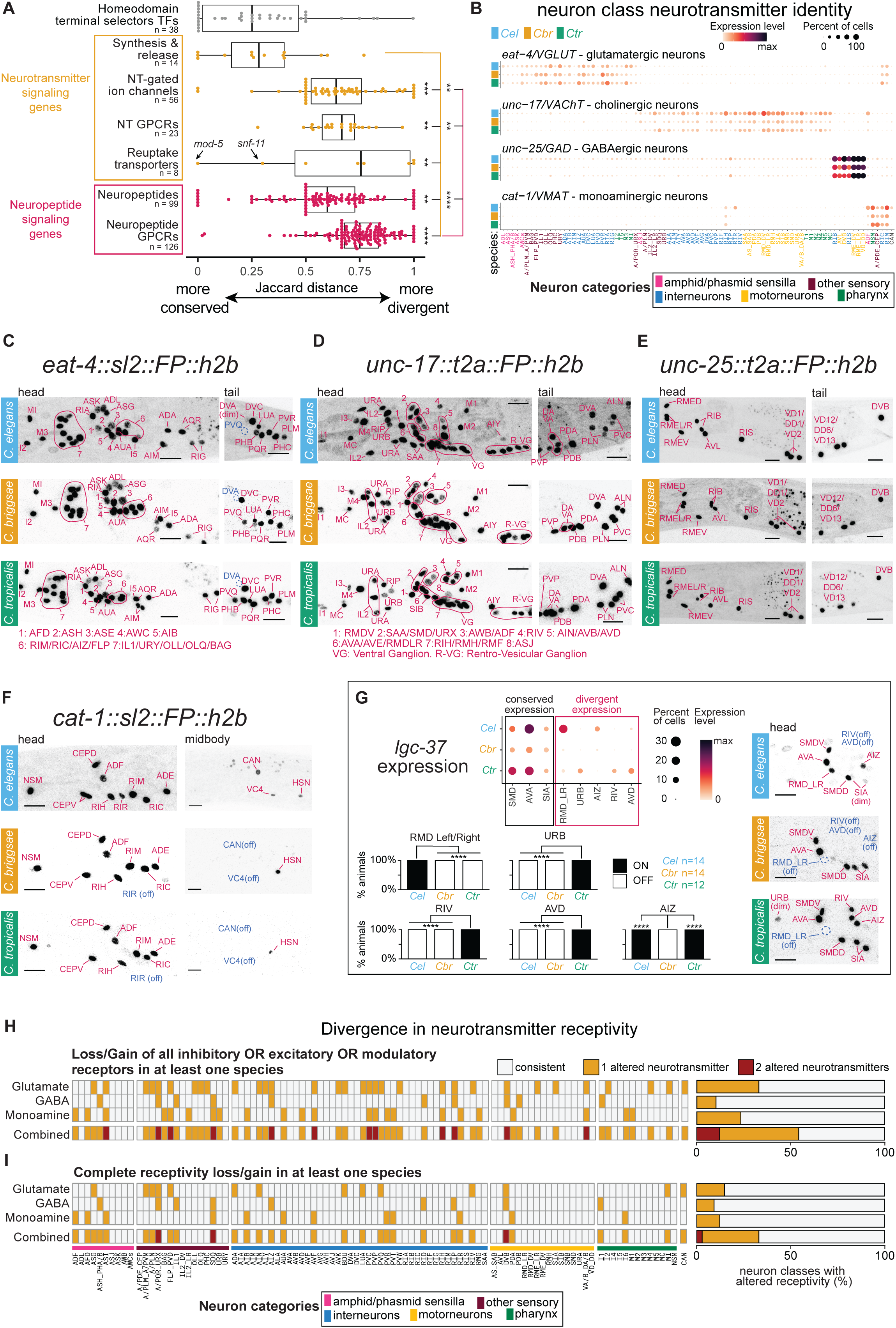
Divergence in cell-type-specific expression of neurotransmitter receptors amid conservation of neurotransmitter release identity. **(A)** Jaccard distances of genes participating in neurotransmitter signaling (yellow) and neuropeptide signaling (red). Each dot represents a gene, boxplots are Tukey-style. n indicates the number of genes present in each gene group. Homeodomain transcription factors data (from Fig. 1I) are shown again for visualization purposes. The neurotransmitter reuptake transporters *mod-5* (serotonin reuptake) and *snf-11* (GABA reuptake), the sole transporters that are essential to confer neurotransmitter identity in some neurons, were the most conserved reuptakers. Kruskal-Wallis test with Dunn’s correction. \*\*\*\**P* < 0.0001, \*\*\**P* < 0.001, \*\**P* < 0.01. **(B)** Cross-species expression dotplot of neurotransmitter synthesis & vesicular transporter genes determining neurotransmitter release identity of neuronal cell classes. Nematode species (y-axis) and neuron class (x-axis) are color-coded according to legend. Neuron classes negative to all 4 genes were excluded. Dot size represents the fraction of cells expressing the gene in a given neuron class, color represents scaled average expression levels. **(C-F)** Representative fluorescent microscopy images (Z-stack max projections, Scale bars: 10μm) of strains with reporter alleles tagging expression of *eat-4/VGLUT* (C)*, unc-17/VAChT* (D)*, unc-25/GAD* (E) and *cat-1/VMAT* (F). Names of expressing neuron classes are indicated in red. VG: Ventral Ganglion. R-VG: Retro-Vesicular Ganglion. Out-of-frame cells with stable signal across species include ALM;AVM;PVD (*eat-4+*), HSN;SDQ;PVN and cholinergic ventral nerve cord motor neurons (*unc-17+*), VD;DD neurons (*unc-25+*), PDE (*cat-1+*). Detectable differences include *eat-4* in DVA neurons (dim in *Cel*, OFF in *Cbr* & *Ctr*), *eat-4* in PVQ neurons (ON in *Cbr* & *Ctr* and OFF in *Cel*), *cat-1* in RIR, CAN, VC4-VC5 neurons (ON in *Cel,* OFF in *Cbr & Ctr*). **(G)** Representative images and quantifications of neuron-class-specific expression of the GABA-gated ion channel gene *lgc-37* in head neurons (Z-stack max projections, Scale bars: 10μm). Sites of expression and divergence are labeled. Corresponding scRNA-seq expression data are shown in dotplots. Barplots represent proportions of scored animals expressing *lgc-37* in the indicated neuron class. Fisher’s test with Bonferroni correction to multiple comparisons. \*\*\*\**P* < 0.0001. The *C. briggsae* and *C. tropicalis* strains contain a CRISPR-inserted ::t2a::mScarlet3::h2b endogenous reporter allele. For the *C. elegans* strain, a fosmid-based construct was used. Cell IDs for RIV, AVD and AIZ were based on cell position and sequencing data, RMD_LR was confirmed using an srlf-7::gfp array. **(H)** Heatmap representing neuron classes in which expression of receptors for the specified neurotransmitter ligand (y-axis) were consistent in all 3 species (grey) or altered (colored). Expression was considered altered if a neuron class expresses ≥2 receptors of a certain type in at least one species and 0 receptors in another species. Inhibitory, excitatory and modulatory receptors for a same ligand were considered separately. Cumulative data is represented in barplots on the right. Full data is shown in Figure S7, information about included receptors are shown in table S5. Several acetylcholine receptors were broadly expressed throughout the nervous system, which meant the strict criteria for receptor divergence was not met by cholinergic receptors. **(I)** As in (H) but all receptors within a neurotransmitter system were considered together. Therefore, a colored neuron class indicates a complete loss/gain of receptivity to the neurotransmitter in one of the nematodes.

To validate this notion, we generated CRISPR/Cas9-engineered knock-in fluorescent reporter alleles of *C. briggsae* and *C. tropicalis* genes that mark the different neurotransmitter systems in nematode, namely *eat-4/VGLUT* (glutamatergic neurons), *unc-17/VAchT* (cholinergic neurons), *unc-25*/*GAD* (GABA-synthesizing neurons), *cat-1/VMAT* (monoaminergic neurons) and compared the expression patterns to the available complete map of neurotransmitter identity in *C. elegans* (Figure 3C-F)(Wang *et al*. 2024). Overall, this reporter analysis confirms conserved neuron-class-specific expression of these three neurotransmitter identity features. We detected subtle cases of species-specific differences in dimly-expressing cells, which we describe in the supplementary text.

In stark contrast to genes conferring neurotransmitter identity to the *sending* neurons, we found that genes encoding for neurotransmitter *receptors* display high divergence in their neuron-class-specific expression across species (Figure 3A). This was true both for neurotransmitter-gated ion channels (n=58 genes with 1:1:1 orthologs) and neurotransmitter metabotropic receptors (n=23) and was independent of neurotransmitter system (Figure S6; table S5). Using CRISPR/Cas9-genome engineered reporter alleles, we validated the species-specific components of the expression patterns of the alpha-subunit type GABA_A_ receptor *lgc-37* in each of the three *Caenorhabditis* species (Figure 3G). In aggregate, we found that 54% of all neuron classes throughout the entire nervous system lost or gained at least one type of neurotransmitter receptivity, defined by the loss/gain of all excitatory, all inhibitory or all modulatory receptors for a given neurotransmitter group (glutamate, GABA, monoamines) in one species but not the others (Figure 3H and S7). Remarkably, 33% of all neuron classes lost all their known receptors to a given neurotransmitter, becoming selectively “neurotransmitter-deaf” in at least one species (Figure 3I and S7). This occurred in 14 neuron types for glutamate receptivity, in 9 neuron types for GABA receptivity and in 12 neuron types for receptivity to all monoamines. These losses are distributed throughout the entire nervous system and show no bias for sensory, inter, motor or enteric neurons.

Taken together, our results indicate a strong selective pressure on cells to retain stable neurotransmitter identity on the releasing side, while displaying rapid divergence as receivers of neurotransmitter-mediated signals.

### Evolutionary plasticity of neuropeptidergic signaling networks

Apart from using one or two classes of neurotransmitters, each *C. elegans* neuron expresses a multitude of neuropeptide-encoding genes, as well as neuropeptide receptors, generating dense “wireless” signaling networks throughout the entire nematode nervous system (Taylor *et al*. 2021; Beets *et al*. 2023; Ripoll-Sanchez *et al*. 2023; Watteyne *et al*. 2024). Our analysis revealed significant divergence in neuropeptide signaling pathways across *Caenorhabditis* species (Figure 3A, *P* < 0.01). In this case, and in contrast to the trend observed in neurotransmitter signaling genes, the divergence was evident on both the signaling and receiving ends. Neuropeptide receptor genes were more divergent in their neuron-type-specific expression than neuropeptide genes (Figure 3A, *P* < 0.0001). We confirmed cases of neuron-type-specific divergence across species with genome-inserted fluorescent reporter alleles expressed in the endogenous regulatory context of the neuropeptide precursor genes. For example, we found that the neuropeptide *nlp-3* (Figure 4A), which is consistently expressed in neurons AWB AWC BAG & NSM, displays divergent expression in ASK (ON only in *C. elegans*), I6 (ON only in *C. briggsae*) I1 I2 & I3 (OFF only in *C. tropicalis*). Likewise, our endogenous *nlp-18* reporter alleles were expressed in *C. elegans* DVB but not DVA, and vice versa in *C. briggsae* & *C. tropicalis* (Figure 4B), and *nlp-11* exhibited divergence in expression in a multitude of neuron classes (Figure 4C).

**Figure 4.**
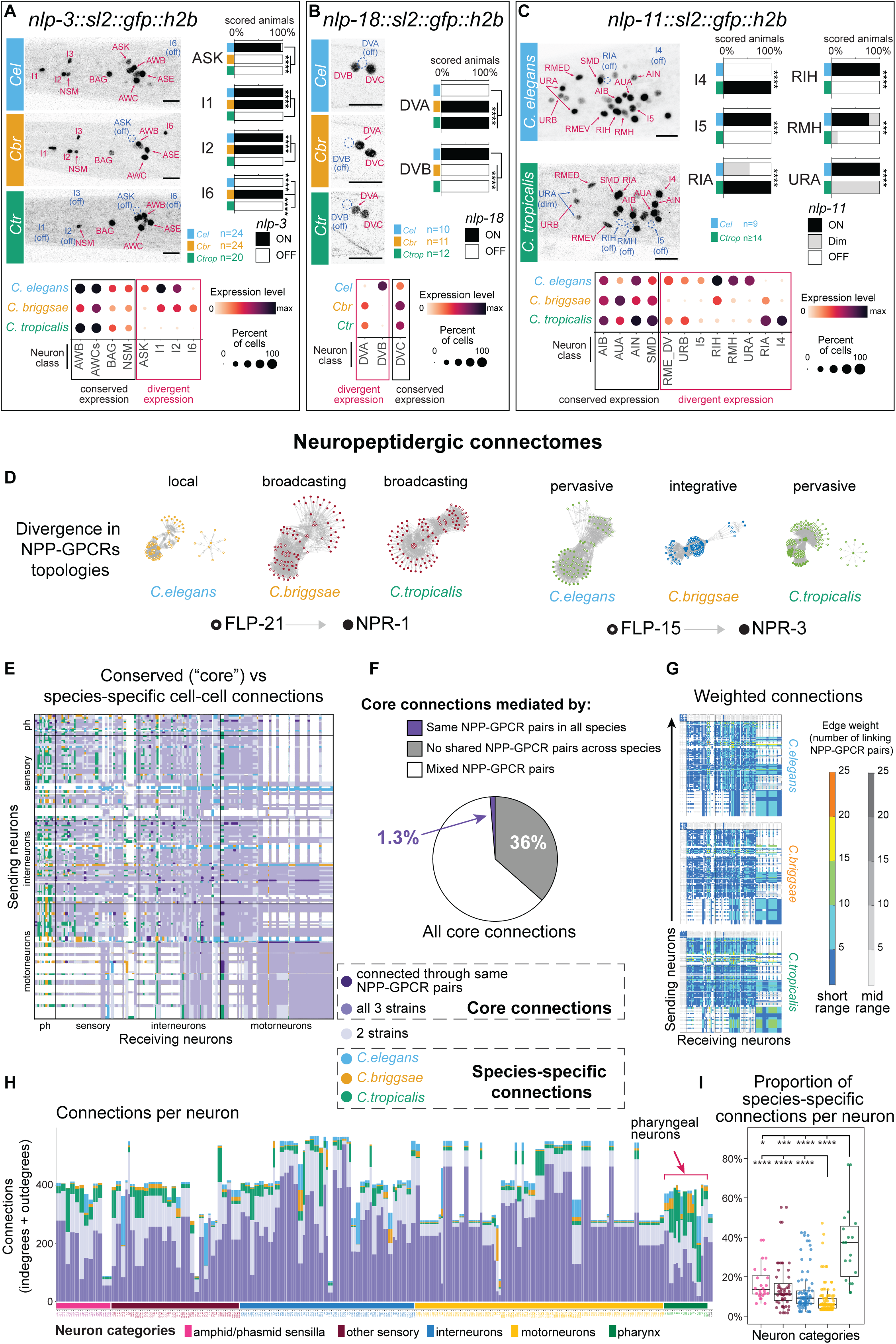
Evolutionary plasticity of neuropeptidergic signaling networks. **(A-C)** Representative images and quantifications of neuron-class-specific expression of neuropeptide precursor genes *nlp-3* (A), *nlp-18* (B) and *nlp-11* (C) tagged with *sl2::gfp::h2b* fluorescent reporter alleles in their endogenous genomic loci (Z-stack max projections, Scale bars: 10μm). Sites of expression and divergence are labeled on the fluorescent microscopy images. Barplots represent proportions of scored animals expressing the corresponding gene in the indicated neuron class. Numbers of scored animals (n) appear below the barplots. Fisher’s test with Bonferroni correction to multiple comparisons. \*\*\*\**P* < 0.0001, \*\*\**P* < 0.001. Corresponding scRNA-seq expression data are shown in dotplots (bottom). (D) Network topologies of the indicated NPP/GPCR pairs. Empty circles represent neuropeptide expression, filled circles represent receptor expression. Local networks display restricted NPP and GPCR expression (≤50 neurons), pervasive display broad NPP and GPCR expression (>50 neurons), broadcaster networks display restricted NPP but broad GPCR expression, integrative networks display broad NPP but restricted GPCR expression. (E) Thresholded neuropeptidergic connectome (mid-range) showing the evolutionary pattern of connections between sending neurons (y-axis) and receiving neurons (x-axis). Core (conserved) connections and species-specific connections are color-coded. To be included, neuropeptide-receptor couples (in their *C. elegans* version) had to pass the functionally validated threshold of EC_50_ < 500nM binding *in vitro* (Beets *et al*. 2023; Ripoll-Sanchez *et al*. 2023). (F) Analysis of the subset of conserved connections producing the core connectome. 1.3% of connections are formed by the same sets of NPP-GPCR pairs across the three species. 36% percent connections share no common NPP-GPCR across the three species. The remaining connections share some but not all NPP-GPCR pairs in common. (G) Weighted neuropeptidergic connectomes, indicating for every cell-cell connection how many pairs of neuropeptide-receptors mediate the connections. (H) Total number of degrees (y-axis, mid-range networks) in homologous neurons across species (x-axis) classified by functional categories. Bars are color-filled according to the subsets of core degrees and species-specific degrees of the neuron across species. (I) Proportions of species-specific degrees (y-axis) per neuron classified by functional categories (colors and x-axis). Kruskal-Wallis test with Dunn’s correction, \*\*\*\**P* < 0.0001, \*\*\**P* < 0.001, \**P* < 0.025.

Next, we assembled “wireless connectomes” to dissect the evolutionary plasticity of neuropeptidergic signal flow in the nervous system (Beets *et al*. 2023; Ripoll-Sanchez *et al*. 2023). We included biochemically validated neuropeptide/receptor pairs, with 1:1:1 orthologs in all three species (42 neuropeptide precursor genes [NPP] and 47 peptide GPCR genes, forming 84 unique ligand-receptor pairs). Analysis of their expression patterns revealed a notable evolutionary divergence of network topologies. Network topologies can be classified into local, integrative, broadcasting or pervasive signaling based on the number of neuron types in which the NPP and the GPCR of a designated couple are expressed (Figure 4D and S8). In 19 (23%) NPP-GPCR couples included in the analysis, cell-type specific changes in gene expression resulted in a topology difference between species (Figure S8 and table S6). For example, expression patterns of the neuropeptide precursor *flp-21* and its biochemically confirmed receptor *npr-1* form a “local” network in *C. elegans* since both genes are expressed in a limited number of cells, while in *C. briggsae* and *C. tropicalis* this pair forms a “broadcasting” network due to extended expression breadth of the receptor *npr-1* (Figure 4D). In another example, the networks formed by *flp-15* and *npr-3* display an integrative topology (many senders and few receivers) in *C. briggsae* but pervasive topologies (many senders and receivers) in *C. elegans* and *C. tropicalis*, Figure 4D).

On a global level, the neuropeptidergic networks of all three species are highly connected (Figure 4E-G). Between 70% and 84% (depending on the inquired species and the network range) of cell-cell wireless connections found in one species were conserved across all three species (Figure 4E; “core connectome”). Yet, intriguingly, the molecular entities that assemble the core wireless connectome (i.e. the particular neuropeptides and GPCR-encoding genes) display widespread drift between species. Indeed, only ∼1.2% (336/28,351 short-range and 476/35,791 mid-range) of the connections in the core connectome are formed via the same sets of NPP-GPCR pairs, and 36% of the core connections share no common NPP-GPCR orthologous pair across the three species (Figure 4E-F).

Further support to the notion of conserved meta-structures of neuropeptidergic communication amid pervasive changes in expression patterns of individual neuropeptidergic genes comes from our analysis of the peptidergic degrees of different neuron classes across the nervous system. Peptidergic degree is defined as the number of incoming and outgoing connections per neuron. Our results indicate very strong cross-species correlations (r = 0.87∼0.9) of degrees between homologous neuron classes (Figure S9). Accordingly, densely-connected “peptidergic hubs” (Taylor *et al*. 2021; Beets *et al*. 2023; Ripoll-Sanchez *et al*. 2023; Watteyne *et al*. 2024) were maintained across species, as well as the general structure of the distributions of degrees in neuronal types (Figure S9 & table S6). This means that despite the pronounced evolutionary divergence in cell-specific expression of neuropeptide and receptor genes, homologous cells tend to retain similar connectivity patterns both as emitting and receiving cells, revelatory of evolutionary pressure acting on the global structure of neuropeptidergic signaling networks.

Alongside the core connectome, each of the species also displays species-specific cell-cell neuropeptidergic connections (Figure 4E), constituting 5% of all connections in *C. elegans*, 2% in *C. briggsae* and 8% in *C. tropicalis*. Close examination of these species-specific edges revealed that they were disproportionally more frequent among cell-cell connections to and from the enteric neurons of the pharynx (Figure 4E, H-I, *P* < 10^-9^ hypergeometric test). The enteric nervous system of the pharynx is synaptically connected to the rest of the central nervous system through a single neuron (RIP), but has previously been appreciated to show extensive communication to the central nervous system via wireless peptidergic connections (Beets *et al*. 2023; Ripoll-Sanchez *et al*. 2023). The species-specificity of both incoming and outgoing connections to the enteric nervous system were proportionally 2.8x∼6x more abundant in pharyngeal neurons than in all other categories of neurons (Figure 4H-I; Figure S9). Together, our findings extend our appreciation for peptidergic signaling operating between the enteric and somatic nervous systems and illustrate that this type of interorgan cross talk is more evolvable than other pathways of communication.

### Novel signaling pathways and their evolutionary divergence

We probed the possible existence and evolutionary plasticity of previously unexplored neuronal signaling pathways by considering the large family of GPCR-encoding genes (1,596 in *C. elegans*, 761 in *C. briggsae*, 1,042 in *C. tropicalis*). We subtracted from these lists GPCRs that are either (a) sequence homologs to known GPCR-type neurotransmitter receptors and neuropeptide receptors, (b) are exclusively expressed in sensory neurons and, hence, likely chemosensory receptors for external cues or (c) are not robustly expressed in any neuronal cell type. This left 51 *C. elegans,* 63 *C. briggsae* and 69 *C. tropicalis* GPCR-encoding genes, all of which candidate receptors for internal signaling molecules (Figure 5A, S10 and table S7). Of those genes, we detected neuron-type specificity (arbitrarily defined here as expression in <20 of neuron classes) in 50/51 (98%) *C. elegans* GPCRs, 57/63 (90%) in *C. briggsae* and 67/69 (97%) in *C. tropicalis.* This apparent specificity in gene expression indicates a possible role in neuron type-specific modulation of information flow. Neurons that expressed the highest number of such GPCRs in all species include the neuropeptidergic hubs PVQ and PVT (Ripoll-Sanchez *et al*. 2023), the octopaminergic class RIC, as well as AIN, AIM, RIP, I5 and CAN (Figure S10). A total of 38 genes are conserved 1:1:1 orthologs and their expression in non-sensory neurons in at least one species leads us to suggest that these receptors may respond to internal signals. Based on Jaccard distance, the expression patterns of these 1:1:1 orthologs are divergent, even more so than the divergence of the neurotransmitter and neuropeptide GPCRs (Figure 5B).

**Figure 5.**
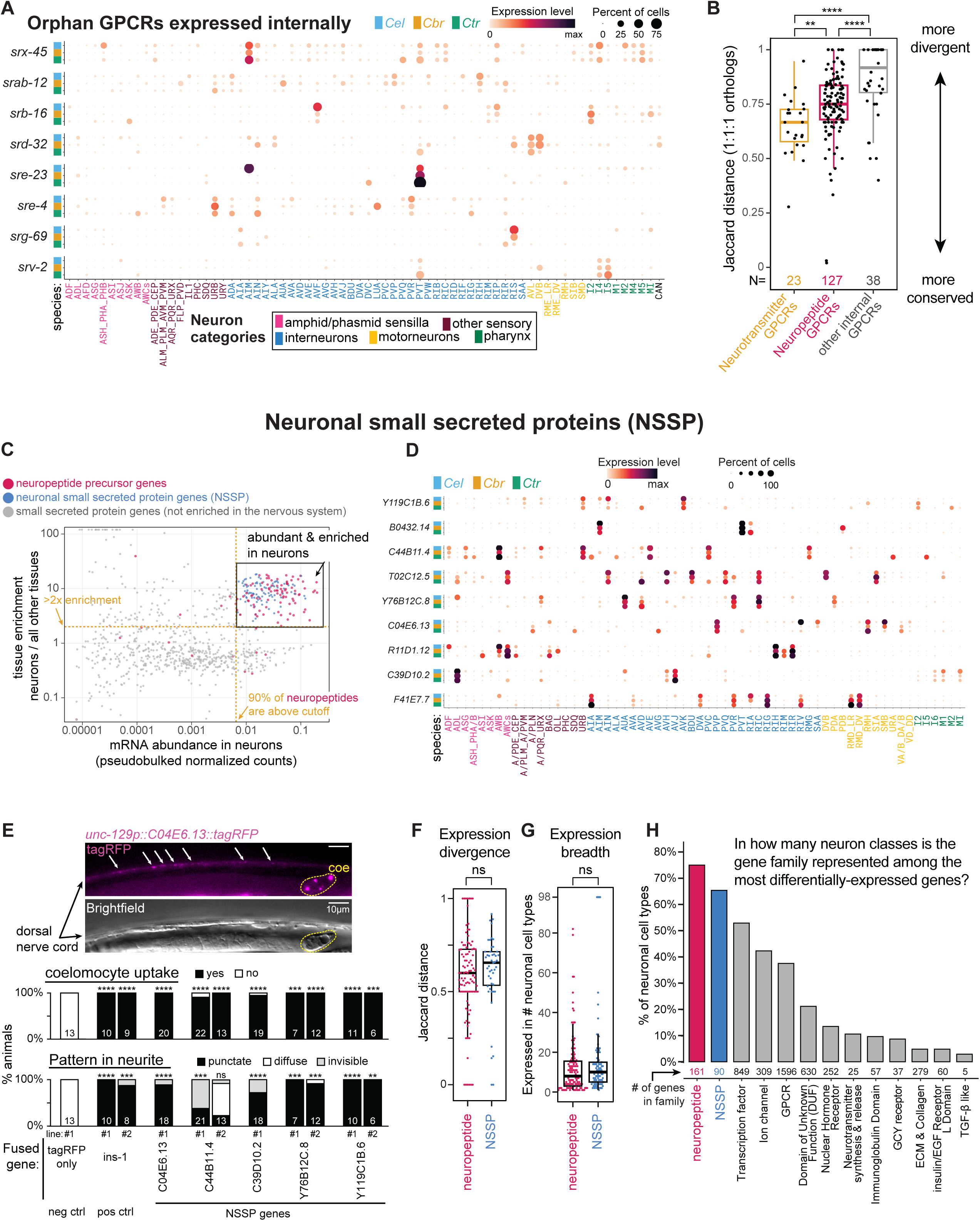
Neuronal expression and evolutionary divergence of gene families potentially participating in novel signaling pathways. (A) Cross-species expression dotplot of a subset of orphan GPCRs that are expressed in non-sensory neurons and have no sequence homology with neuropeptide- and - neurotransmitter-binding GPCRs. Nematode species (y-axis) and neuron class (x-axis) are color-coded according to legend. Dot size represents the fraction of cells expressing the gene in a given neuron class, color represents scaled average expression levels. (B) Jaccard distances of 1:1:1 GPCR orthologs, grouped according to their known ligand. Each dot represents a gene, boxplots are Tukey-style. N indicates the number of genes per group. Kruskal-Wallis test with Dunn’s correction. \*\*\*\**P* < 0.0001, \*\**P* < 0.01. (C) Neuronal enrichment (y-axis) and neuronal expression levels (x-axis) of genes encoding small secreted proteins in *C. elegans*. Each dot is a gene, neuropeptides (red) and the novel family of NSSPs (blue) are colored. Dashed lines: cutoff criteria used to delineate NSSPs. (D) Cross-species expression dotplot of a subset of conserved 1:1:1 NSSP orthologs. NSSPs tend to be expressed at high levels in few neuron classes. (E) Expression of NSSP::tagRFP protein fusion transgenes in cholinergic motor neurons (unc-129 promoter). Secreted proteins are uptaken by coelomocytes, resulting in detectable fluorescence therein. Localization in axonal release vesicles can be observed by punctate expression in the dorsal nerve cord neurites. Top – representative images of unc-129p::C04E6.13::tagRFP fluorescence pattern (magenta). Coelomocyte is marked in yellow. Arrows mark punctate fluorescence in the nerve cord. Scale bar - 10μm. Bottom – Quantifications of coelomocytes uptake and expression pattern in nerve cord (Percentage out of all tested animals, y-axis). Fused protein shown in x-axis. unc-129p::tagRFP with no fused protein was used as negative control. unc-129p::INS-1::tagRFP was used as positive control. N of total tested animals shown inside barplots. Fisher’s exact test with pairwise comparisons to negative control and Bonferroni correction, \*\**P <* 0.01, *\*\*\*P* < 0.001, *\*\*\*\*P* < 0.0001. (F) Jaccard distances of 1:1:1 neuropeptides and NSSPs ortholog genes. Wilcoxon rank sum test, *P* > 0.05. (G) Expression breadth of neuropeptides and NSSPs in *C. elegans*. Wilcoxon rank sum test, *P* > 0.05. (H) Proportions (y-axis) of neuron classes that include genes from indicated gene family (x-axis) among their most differentially-expressed genes (top_markers score > 0.1) in *C. elegans*. Number of all genes belonging to each gene family (not only the differentially-expressed genes) appears below bars.

Since GPCRs are well known to bind small peptides (Beets *et al*. 2023), we explored the three *Caenorhabditis* genomes and expression datasets for small secreted “orphan” peptides, that may participate in yet undetermined modes of neuronal signaling. To delineate a pool of “<u>n</u>euronal <u>s</u>mall <u>s</u>ecreted <u>p</u>roteins” (from here on referred to as “NSSPs”) in each nematode species, we first filtered predicted proteins shorter than 200 amino acids (>95% of known neuropeptide precursor proteins in *C. elegans* are shorter than this cutoff, Figure S11). We further filtered the genes based on the presence of a signal peptide, lack of transmembrane domains, lack of predicted enzymatic domains and gene expression features (tissue-specificity and mRNA abundance) (Figure 5C, S11, table S8). These criteria were fulfilled by 90 *(C. elegans*), 84 (*C. briggsae*) and 88 (*C. tropicalis*) NSSPs, including 47 (52%∼56%) 1:1:1 orthologous NSSPs expressed in neurons of all three species (Figure 5C-D, S11-12, table S8). None of these genes were previously described or characterized. We expressed 5 conserved NSSPs fused to tagRFP in motor neurons of the ventral nerve cord (Figure 5E) and observed protein uptake by scavenger-like coelomocytes (5/5) and punctate localization at presumptive axonal release sites (4/5), both features of secretion that are displayed by canonical neuropeptides (Sieburth *et al*. 2005; Sieburth *et al*. 2007; Laurent *et al*. 2018).

Given the way we filtered and delineated the pools of NSSPs, it is unsurprising that these genes share with neuropeptides genes the feature of highly abundant expression levels in the nervous system. Unexpectedly however, we found that NSSPs and neuropeptides shared additional expression and evolutionary features in common. Neuron-class specific expression of neuropeptide and NSSP 1:1:1 ortholog genes exhibited similar rates of evolutionary divergence, as reflected in their Jaccard distances (Figure 5F). Genes from both groups also tend to be sparsely expressed in a limited number of neuronal cell classes, that in aggregate span the entire nervous system (Figure 5D & G and S12-13). Like neuropeptides, NSSPs are overrepresented among the most abundantly and class-specifically expressed genes in the nervous system (Figure 5H and S12), making them a key distinctive molecular feature of a majority of neuron classes. Perhaps most remarkably, like neuropeptides, each neuron class expressed a unique combination of multiple NSSP genes (averaging 14 genes). In conclusion, NSSPs, perhaps via GPCR-type receptor, may define previously unappreciated signaling axes in nematode nervous systems.

## Discussion

We have leveraged here the compact nature of nematode nervous systems and their well-annotated genomes to reveal patterns of evolutionary changes throughout the entire brains of species that diverged >40 millions of years ago. We found no evidence for the evolution of novel neuronal cell types; rather each individual neuron class could be readily homologized by either considering the entire battery of genes expressed in a neuron type or, more succinctly, by the combination of homeodomain transcription factors. It has been previously appreciated that transcription factors provide a key proxy for cell type classification (Arendt *et al*. 2016; Arendt *et al*. 2019). Our deep knowledge of transcription factor function in *C. elegans*, developed over several decades of genetic loss of function studies (Hobert 2016), reveals that not all of the many transcription factors that are expressed in a mature neuron class carry similar weight for proper neuronal classification. It has rather become clear, and further validated here in the context of additional species, that of all transcription factor families, homeobox genes represent the most potent classifiers of neuronal identity (Reilly *et al*. 2020). This insight may help to classify neuronal cell types in other organisms, particularly in those with limited available data on transcription factor function.

Aside from the apparent stability of at least specific subsets of transcription factors, our analysis describes a wide range of molecular changes in nematode nervous systems. The nervous system-wide nature of our analysis allowed us to ask whether specific parts of the nervous system evolve more rapidly than others. Accumulating evidence from diverse phyla suggested that the sensory apparatus is more evolutionary labile than other neuronal features, a reflection of successful adaptations to varying ecological and behavioral niches with different sensory requirements (Bendesky and Bargmann 2011; Mcgrath *et al*. 2011; Cande *et al*. 2013; Auer *et al*. 2020; Roberts *et al*. 2022; Ma and Zheng 2023; Ma *et al*. 2024). We found that sensory neurons indeed diverge more rapidly in the expression of non-conserved genes. Interestingly, these changes are not solely the result of gain or loss of sensory receptor genes but encompass many genes with no known protein domains. However, sensory neurons do not stand out in the context of evolving new expression of conserved genes. Unexpectedly, we found that motor neurons differ from all other types of neurons in their stability of expression of orthologous genes, hinting towards constraints in the evolvability of the last step of information flow from the nervous system to motor end organs.

Our analysis of neuronal signaling pathways in the nervous system has revealed striking patterns of evolutionary divergence. While the deployment of a given classic neurotransmitter system is remarkably stable over evolutionary time and neurotransmitter type (ACh, GABA, Glu, monoamines), the patterns of reception of such signal display dramatic changes, as inferred by highly plastic expression patterns of all types of neurotransmitter receptors. These changes are best illustrated in the “broadcasting” network structure of monoaminergic signals (Bentley *et al*. 2016) where a very small number of either serotonergic, dopaminergic, tyraminergic and octopaminergic neurons, respectively, retain their signaling capacity, but a distinct set of downstream neurons listens to these inputs in different nematode species. Taken together, in spite of an overall conserved neuronal architecture, signal flow through classic neurotransmitter systems is highly evolvable.

In addition to their use of classic neurotransmitter systems, nematode neurons express a combination of around 20 neuropeptide-encoding genes per neuron class (Taylor *et al*. 2021). We found that the expression of individual neuropeptide precursors and cognate receptor pairs is also rapidly evolving. However, thanks to our whole-nervous-system and single-neuron perspective, we discovered that despite widespread divergences at the level of individual genes, the vast majority of putative neuropeptidergic connections between pairs of homologous neurons were conserved across species. This observation may indicate stabilizing selection that maintains, in diverging species, adaptive context-dependent neuromodulatory states through compensatory routes of cell-cell interconnections. “Re-coding” of pathways of neuropeptidergic communication through compensatory gain and losses of neuropeptide and receptor expression is enabled by the deep repertoire of neuropeptides and neuropeptide receptors expressed by each individual neuron (Taylor et al. 2021).

While compensatory changes make the neuropeptidergic connectome robust to changes in neuropeptidergic information flow in the central nervous system, the enteric nervous system of nematodes, located in the foregut (pharynx), stood out as a hotspot for rapid evolutionary divergence in its peptidergic connectivity. This was reflected in the disproportionately high concentration of species-specific edges connecting pharyngeal neurons with each other or connecting pharyngeal with somatic neurons, in both directions. Throughout animal phylogeny, enteric nervous systems often form self-contained and largely (but not completely) autonomous circuits (Copenhaver 2007; Sasselli *et al*. 2012). Animal enteric nervous systems not only control proper digestive activity, but are also being recognized as key sensory hubs that perceive a wide range of signals from food, ranging from nutritiousness to pathogenicity (Furness *et al*. 2013; Yoo and Mazmanian 2017). Species-specific differences in cellular inter-connections might reflect functional adaptations to distinct ecological demands acting on the coordination between the enteric and somatic nervous systems to tune organismal responses to nutrients or pathogens.

Our comparative mining of the three nematode brain atlases also provides tantalizing hints for the existence of neuronal signaling modules that go beyond canonical neurotransmitter and neuropeptide signaling pathways. We described a multitude of GPCR-encoding genes, as well as small, secreted peptides, which show very selective patterns of expression with the brains of the three nematodes. The GPCRs may be receptors for internally secreted metabolites, for example ascarosides (Ludewig and Schroeder 2013), or may find their ligand among those neuronally small secreted proteins (NSSP) that we described. Our observation of each individual neuron class expressing a unique combination of, on average, 14 NSSPs genes almost doubles the already very impressive repertoire of peptidergic signals emanating from neurons. GPCRs and NSSPs that are conserved across the three nematode species show patterns of divergence in their neuronal expression that are similar to those seen for neuropeptides and their receptors, if not even more pronounced. Whether these putative signaling systems may be drivers of behavioral changes remains to be investigated, but in any case, their mere existence points to presently perhaps underappreciated depth of signaling pathways within the nervous system.

## Methods

### Cultivation of nematodes for scRNA-seq experiments

*C. elegans* (strain N2), *C. briggsae* (AF16) and *C. tropicalis* (NIC203) were cultured at 20°C with *E. coli* strain OP50 using standard conditions with the exception that the agar in the nematode growth media (NGM) was replaced with a 4:6 mixture of agarose and agar (NGM+agarose) to prevent burrowing. To generate a large synchronized population of L2 worms for single-cell RNA-seq (scRNA-seq) experiments, adult hermaphrodites from the three strains were treated with hypochlorite solution and the resulting embryos were kept overnight (∼16h) in M9 buffer to hatch and arrest in L1 stage. Then, L1s were transferred to 10cm plates pre-seeded with OP50 (4 plates, 50,000 worms/plate, total of 200,000 worms per strain). After 23 hours, the L2 stage was verified under a stereoscope.

### Cell dissociation for single-cell RNA-sequencing

Cell dissociation was carried out as previously described (Ben-David *et al*. 2021a) with minor modifications for the three worm species. L2 worms were recovered off the plates and washed three times in M9 in a 15ml conical tube, followed by two times in a 1.5ml tube. Lysis was then performed using a freshly thawed aliquot of 200μl SDS-DTT solution (200 mM DTT, 0.25% SDS, 20 mM HEPES, pH 8.0, 3% sucrose) for 6 minutes in a hula mixer set on low speed. Worms were then washed quickly three times in 1ml of M9, and two additional times in 1ml of egg buffer (118 mM NaCl, 48 mM KCl, 2 mM CaCl2, 2 mM MgCl2, 25 mM HEPES, pH 7.3, osmolarity adjusted to 340 mOsm with sucrose). Worms were then resuspended in 500μl of 20 mg/ml Pronase E (Sigma-Aldrich, P8811) that was freshly prepared in L15-FBS (L15 medium supplanted with 2% fetal bovine serum and adjusted to 340 mOsm with sucrose). Worm dissociation was done by continuous pipetting on the side of the tube. The dissociation process was monitored every 2-3 min on a microscope equipped with a x40 phase contrast objective lens. Dissociation was stopped when a high density of cells was visible. Dissociation durations were different for each species: 8:36 for *C. elegans*, 13:16 for *C. tropicalis*, and 17:16 for *C. briggsae*. Dissociations were stopped by adding 500μl ice-cold L15-FBS and transferring the tubes on ice. Following dissociation, lysates were spun for 6 min at 500g at 4°C. Cell pellets were resuspended in cold PBS (adjusted to 340 mOsm with sucrose). Cell suspensions were spun for 1 min in 50g to pellet remaining undigested worms. Cell preparations (supernatants) were transferred to new 1.5ml tubes (pre-cooled on ice) and cells were counted on a hemocytometer loaded to an inverted microscope equipped with differential interference contrast (DIC). Cells from each species were then diluted to 10^6^ cells/ml concentration in osmolarity-adjusted PBS, and then combined together. The combined pool was loaded onto eight lanes of 3′ Chromium scRNA-seq flow cells (10x Genomics), targeting 30,000 cells on each lane. Library prep was carried out according to the manufacturer’s protocol. Libraries were sequenced together on three S4 lanes of Novaseq 6000. Paired-end 2 × 150 runs were done to maximize the recovery of transcript variants between the three species.

### Genome engineering and transgenics

Knock-in reporter alleles were generated using CRISPR/Cas9 or Cas12a and single-stranded oligodeoxynucleotides (ssODNs) for precise insertions. Injection mixtures were prepared using enzymes and RNAs ordered from IDT (Cas9 #1081059, L.b.Cas12a #10007922, tracrRNA #1072532) and according to the injection procedure and concentrations described in (Ghanta and Mello 2020). Cas9 (0.5μl of 10μg/μl stock), tracrRNA (5μl of 0.4 μg/μl stock) and crRNA (2.8μl of 0.4 μg/μl stock) were gently mixed together and left to incubate at 37°C for 15 minutes to form the RNP complex. The ssODN (2.2μg) was then added and mixture complemented with nuclease-free water to a final volume of 20μl, used for microinjection. F1 progeny of injected hermaphrodites were screened for engineered heterozygotes through examination under a fluorescent dissection microscope (Leica M165FC) and PCR genotyping, before isolation of homozygotes in subsequent generations. All insertions in the resulting strains were validated by Sanger sequencing.

The ssODNs used as CRISPR repair donor templates were relatively long (typically 0.9kb∼1.6kb). They all included the desired insertion sequence (including mutations in the PAM or crRNA complementary region when needed) flanked by 35bp-homology arms. Long ssODNs were prepared using a previously-described procedure based on the enzymatic digestion of PCR products into single-stranded DNA (Eroglu *et al*. 2023). We designed PCRs amplifying exactly the desired repair template sequence with one (but not both) primer bearing a phosphorylated 5’ nucleotide. PCRs were run in 50μl reactions (4 side-by-side replicates) using Q5 enzyme and buffers (NEB; M0491). Amplification was confirmed through agarose gel electrophoresis, then the replicates were pooled together and column-purified (Invitrogen^TM^ PureLink^TM^ #K310001). Eluted purified PCR products (45μl in nuclease-free water) were digested with a lambda exonuclease (NEB M0262L) and accompanying reaction buffer for 20 minutes at 37°C (total reaction volume 50μl). Lambda exonuclease is a strand-specific exonuclease that preferentially degrades DNA strands phosphorylated at their 5’ end. Enzymatic digestion was followed by column purification using the Monarch® system (NEB T3010S) and elution with 6μl of nuclease-free water.

Transgenic strains expressing NSSP genes fused to tagRFP in the motor neurons of the ventral nerve cord were obtained by co-injecting the NSSP expression plasmid (50ng/μl) and pha-1(+) rescue construct (50ng/μl) into *pha-1(e2123) C. elegans* worms and growing them at 25°C (lethal temperature for non-rescued animals). The structure of all NSSP transgenes was: *unc-129p::NSSP::tagRFP::sl2::gfp::h2b*. An *ins-1* fusion construct was used as positive control (established secreted neuropeptide) and a non-fused *tagRFP* cassette as negative control. Information about strains, crRNAs, ssODNs and plasmids used for this study are found in table S9.

### Microscope Imaging and analysis

Animals were anesthetized in a drop of M9 with 50mM sodium azide and mounted on glass slides padded with a patch of 5% agarose. Images of strains bearing fluorescent endogenous reporter alleles were acquired using Zeiss confocal microscopes LSM880 or LSM980 with a 40x water objective. FIJI software (Schindelin *et al*. 2012) was used to analyze neuron-specific expression and to generate orthogonal Z max projections of representative images. To facilitate neuron identification, animals were stained with Vybrant^TM^ DiD cell-labeling solution (Invitrogen # V22887) and/or crossed with strains expressing fluorescently-tagged genes in conserved sets of cells. Images of NSSP::tagRFP transgenic worms and controls were acquired using a Zeiss Axio Imager Z2 microscope at x40 objective. Images were analyzed to characterize uptake of TagRFP in coelomocytes and for punctate/dispersed expression in the dorsal nerve cord. *nlp-18::gfp* animals were imaged as L2s, NSSP::tagRFP transgenic nematodes were imaged as L4s or young adults, all other images were taken using L4 nematodes.

### Genome builds and *In-silico* demultiplexing for species-specific fastq files and cellranger analysis

The raw sequencing data included reads originating from all three species. To generate species-specific fastq files, we first aligned the reads separately to the *C. elegans* genome (WBcel235)(Sternberg *et al*. 2024), the *C. tropicalis* NIC203 genome (Ben-David *et al*. 2021b), and the recently published *C. briggsae* QX1410 genome (Stevens *et al*. 2022) that was annotated *de-novo* using Funannotate V1.8.7 (https://github.com/nextgenusfs/funannotate/). The longest mRNA isoform of each gene was used for alignment. The gene transfer files (GTF) annotation of 3’UTR regions in *C. briggsae and C. tropicalis* were then dynamically extended to improve the mapping of 3’ scRNA-seq reads, following the same iterative approach outlined in a previous study for *C. elegans* (Packer *et al*. 2019). The alignment to the different species was compared based on the CIGAR string in the alignment bam file, and reads that supported one of the species better than the two others were used to generate three species-specific sets of fastq files. Alignment and counting were then performed using Cellranger v7.0.1 (10x Genomics).

### Identification of gene ortholog sets

To identify gene orthologs, we took advantage of the broad synteny previously reported between *C. elegans* and each of the other two species (Stein *et al*. 2003; Ben-David *et al*. 2021b). We carried out a tiered analysis that considered multiple possible scenarios for orthologous genes and ranked them by the strength of the evidence. Our analysis included both synteny-agnostic orthology discovery with Orthofinder (Emms and Kelly 2019) and synteny-aware orthology discovery using pSONIC (Conover *et al*. 2021). We carried out ortholog gene discovery with each of the tools using either the transcriptome (cDNA) sequences or the proteome (amino acid) sequences. Evidence for orthology was ranked based on the source using the following encoding:

1. Unanimous orthologs, identified unequivocally with a 1:1 mapping across all analyses.
2. Syntenic orthologs discovered at the cDNA level
3. Syntenic orthologs discovered at the protein level
4. Non-syntenic orthologs discovered at the cDNA level
5. Non-syntenic orthologs discovered at the protein level

We then used this annotation list to call a consensus set of 1:1 orthologs using the following algorithm: For every gene (A) in every species, (A) was considered a <u>consensus</u> <u>1:1 ortholog</u> of a gene (A’) in another species if (A) was found homologous only to (A’) when considering the homology levels in decreasing order (i.e. if [A] was found uniquely homologous to [A’] in level 2, consideration was stopped and did not continue to more lenient level 3), and if no other gene (B) was called a consensus 1:1 ortholog to the same (A’) gene in a more stringent homology level. The final set of 1:1:1 ortholog gene triplets (across all three species) included 11,277 triplets.

### Classification of genes with no consensual 1:1 orthologs

Genes that had no 1:1 ortholog genes whatsoever in the two other species in all homology levels tested were considered “1-to-none” novel genes. The “permissive sets” of genes (Figures 2B-C) included all genes that had no <u>consensual</u> 1:1 orthologs in neither of the two other species. Hence, the permissive sets included 1-to-none novel genes as well as genes in orthogroups that include recently-duplicated paralogs or multiple closely-related paralogs in both sequence and synteny. Genes that had a <u>consensual</u> 1:1 ortholog in a second but not a third species were left out of the permissive sets.

### Downstream processing and annotation of cell types

A) *C. elegans* cells differ widely in the number of <u>U</u>nique <u>M</u>olecular Identifiers (UMIs) that are recovered in scRNA-seq. As a result, a simple UMI cutoff may be biased for cell types with more UMIs (Packer *et al*. 2019). Therefore, we implemented an iterative pipeline to recover clusters of high-quality cells and remove cell doublets and degraded cells. For each species, we took 25,000 cells with most UMIs in each lane (2.5x the targeted number of cells, 200,000 overall) and processed them in *Monocle3*(v1.3.4) available at https://cole-trapnell-lab.github.io/monocle3/ (Trapnell *et al*. 2014; Qiu *et al*. 2017; Cao *et al*. 2019). We used default parameters for processing (preprocessing, dimensionality reduction and clustering) except that 100 dimensions were included for PCA. We used the *top_markers* function in *Monocle3* to identify enriched genes in the obtained clusters (Leiden clustering), and removed clusters whose top enriched genes were mostly ribosomal or mitochondrial genes and no known cell type markers. We repeated this cell filtering step a second time with the remaining cells. We then used scrublet(v0.2.3) (Wolock et al. 2019) to remove suspected cell doublets (doublet score > 0.1) and SoupX(v1.6.2) (Young and Behjati 2020) to correct for background contamination by ambient RNAs, reprocessing the datasets after each step. Resulting datasets were used for manual cell type annotations. During downstream analyses, we noticed several cell types missing in our data due to low UMI counts of *Caenorhabditis* neurons, as previously reported (Packer *et al*. 2019). To recover those cells, we used markers co-expressed very specifically in individual neurons to identify them in the raw expression matrix and then reintegrated those cells into the filtered datasets (“reincluded” column in the annotated datasets). The full datasets, including the reintegrated cells, were reprocessed, clustered, corrected for doublets and ambient RNAs before final cell annotations and analyses.

To annotate cell types, we compared the enriched differential genes in each cluster to markers confirmed in previous *C. elegans* studies (Packer *et al*. 2019; Taylor *et al*. 2021), or their 1:1 orthologs for *C. briggsae* and *C. tropicalis*. After assigning clusters into broad tissue types, all cells of a given tissue (for example all neurons) were further subclustered in isolation to separate refined clusters and cell types. Most neuronal clusters could be readily assigned to an individual neuron class or subclass. In other cases, we applied more than one cycle of separated subclustering. When clusters appeared to contain cells from several closely-related neuron classes that couldn’t be confidently separated (such as the oxygen sensing neurons AQR/PQR/URX, or the dopaminergic neurons ADE/PDE/CEP) we annotated them accordingly as a group. In some cases, the granular separation of neuron classes was of higher resolution in one species compared to others. For example, the chemorepulsive neurons ASH, PHA and PHB clustered separately in the *C. elegans* dataset but could not be confidently separated in the other species. Therefore, in the count matrices metadata of each species, we generate distinct columns for the highest-resolution annotations in each species (“within_species” column, e.g. separate ASH, PHA & PHB) and annotations that regrouped cells into common annotations across species (“cross_species” column, e.g. ASH_PHA_PHB). Species-specific analyses (for example in Figure 2A-C) relied on the former, while cross-species comparisons (the majority of analyses presented) relied on the latter. Cross-species comparisons also excluded neuron classes that included no or only very few sequenced cells in the dataset of one of the species (*C. tropicalis* ASE & I3, *C. briggsae* AWA). Lists of top differential genes (*top_markers* outputs) are available in table S2.

### Transcriptomic correlation heatmaps

In each species, we pseudobulked the normalized expression values calculated from *Monocle3* for all genes in every neuron class (cross_species neuron annotations). Then, we collected all genes with 1:1:1 orthologs that had a top_markers score > 0.1 in at least one neuron class and in at least one species (1,380 1:1:1 orthologs in total). The pheatmap(v.1.0.12) function (Kolde 2015) on pearson correlations of pseudocount normalized values were used for hierarchical clustering and visualizations.

### Expression thresholding and calculation of Jaccard distances

To threshold expression data into binary ON/OFF expression values, we trained a random forest classifier on ground truth data from *C. elegans*. Our ground truth expression matrix was based on the dataset previously generated by the CeNGEN project (Taylor *et al*. 2021) which itself had been compiled based experimentally-validated fluorescent reporters. The original ground truth matrix was modified to account for differences in the grouping of neuronal classes and for recent updates in characterizations of expression patterns (Wang *et al*. 2024). To train the random forest model, we included features at the cell-cluster level and at the gene level. At the cell-cluster level, our features included the following matrices:

A) A z-score standardized matrix of normalized expression values that was binarized to include 1 in cells with z>1.5
B) For each gene, the fraction of expressing cells (defined as having raw counts > 0) in each cell type.
C) For each gene, the number of expressing cells (defined as having raw counts > 0) in each cell type.
D) The pseudobulked count matrix
E) For each gene in each cell type, the fraction of cells with counts > 0. That matrix was further normalized by the maximum fraction observed for each gene (“percentile thresholding” in (Taylor *et al*. 2021)).

Gene level features:

F) The variance of the pseudobulked expression matrix
G) The variance of the pseudobulked normalized expression matrix
H) The total number of cells with expression in the gene.

For training the classifier, the genes were split to Training+Validation/Test sets (75%,25%). We then upscaled the positive samples to account for 69%/31% split in the training data. We trained a random forest classifier with 1000 trees using 10-fold cross validation regime on the training data. On the test set of unseen genes with ground truth knowledge in *C. elegans*, our classifier achieved 96% precision and 88% recall. The classifier was used to infer binarized expression in all genes in all three species.

Thresholded expression data of 1:1:1 orthologs was aggregated across species and used to calculate Jaccard distances according to the formula 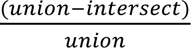. More explicitly:

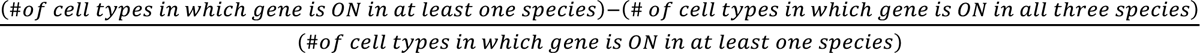

Cell-centered Jaccard distances for each cell type were calculated as:

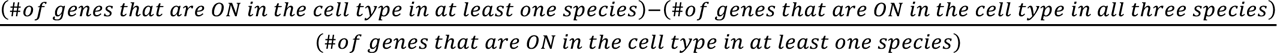

Genes included for the cell-centered Jaccard distances were either all 1:1:1 orthologs expressed anywhere in the nervous system (at least one neuron in any of the three species 9666 genes) or the subset of 1:1:1 orthologs belonging to gene families with established roles in neuronal activity (884 genes, see (Hobert 2013)).

### Classification of genes into gene families

(A) *C. elegans* genes and their orthologs in *C. briggsae* and *C. tropicalis* were assigned to neuronal gene families of interest based on previous research in *C. elegans*. In addition, we used the InterProScan pipeline (https://www.ebi.ac.uk/interpro/) to annotate protein domains based on the predicted translated sequences of all three species using PFAM and the PFAM-A database. We then applied a custom-made pipeline to assign genes to gene families based on family-defining protein domains. Tables for all genes and their respective orthologs, protein domains and annotated families are available in table S9.

### Tree building

To generate schematic phylogenetic trees of gene orthogroups, we first run multiple-sequence alignment (T-Coffee v.11.00) using protein sequences. The MSA output was used to generate trees at phylogeny.fr (Dereeper *et al*. 2008) with default parameters.

### Analysis of evolutionary divergence of neurotransmitter receptors

Lists and details about neurotransmitter receptors in *C. elegans* are found in table S5. Genes with 1:1:1 orthologs were selected and classified according to their category (excitatory ionotropic receptors, inhibitory ionotropic receptors, modulatory metabotropic receptors) and their neurotransmitter ligand (glutamate, GABA, biogenic monoamines). Acetylcholine receptors were left out because some of them (such as *unc-38*, *unc-63* & *acr-12*, shown in Figure S7) were found to be expressed very broadly across the nervous system, mostly at low expression levels, which makes our divergence analysis hard to conclusively interpret for these cases. Thresholded data was used to quantify how many receptors of each category are expressed in each neuron class and species. Receptivity was considered altered across species in a ligand/activity category if at least one species expresses at least 2 receptors in the category and one species expresses 0 receptors in the category.

### Neuropeptidergic connectomes and network analyses

#### Neuropeptide network spatial constraining

Neuropeptidergic signaling was locally thresholded to filter out connections between neurons that were anatomically distant from each other, based on electron microscopy data and neuronal reporter strains of the *C. elegans* nervous system in L2 stage (23h after hatching) (Smith *et al*. 2010; Witvliet *et al*. 2021; Mulcahy *et al*. 2022; Rashid *et al*. 2022). These data were used to create a matrix of locations and anatomical proximity for the processes of each neuron, identifying 27 different neuronal process bundles in the *Caenorhabditis* nervous system as previously defined (White *et al*. 1986). This classification was then used to filter out neuropeptidergic connections based on putative signaling ranges in the three species. The stringent short-range thresholding allows connections only between neuronal processes that are in the same process bundle, and the pharynx is a separated system where connections are allowed between pharyngeal neurons only. The mid-range stringency thresholding allows connections between neurons with neuronal processes in the same anatomical area: head (including pharynx and the ventral cord neurons that are in the ventral ganglion), midbody and tail.

#### Neuropeptide network construction

A previous study biochemically validated interactions between *C. elegans* neuropeptide ligands and GPCR receptors (Beets *et al*. 2023). Out of these validated interaction pairs, we used thresholded expression data for the 42 neuropeptide precursor genes (NPP) and 47 GPCR receptors that had 1:1:1 orthologs in our datasets. Adjacency matrices were built using a binary version of the expression data for the 285 single neurons present in the datasets of the three species. For a given point *A*(*i*, *j*)*^N^* and for a given NPP–GPCR pair N the connection between two neurons is defined by *A*(*i*, *j*)*^N^* = *NPP*(*i*, *j*)*^N^* × *GPCR*(*i*, *j*)*^N^*. Each NPP-GPCR interaction forms an individual binary network. To generate global neuropeptide networks, we summed each individual NPP-GPCR network resulting in weighted networks in which the weight indicates the number of NPP-GPCR pairs that connect two nodes (cells). Reciprocal connections between nodes were considered as two separate unidirectional connections. For the cross-species network analysis, we binarized all the connections (ON if weight of connection ≥1) of each species’ network separately, then integrated the data into a single network reflecting the cross-species conservation pattern of connections between homologous neurons.

#### Topological network measures: degree

Edge counts and adjacency matrices were all computed using binary directed versions of the networks. The same networks were used to compute degree using the method from the Brain Connectivity Toolbox (Rubinov and Sporns 2010) for MATLAB. Degree is the number of edges connected to a given node. In-degree is the number of incoming connections connected to a given node and out-degree is the number of outgoing connections.

### Identification of orphan GPCRs expressed in non-sensory neurons

Lists of all GPCR genes in each species were filtered to remove neurotransmitter- and neuropeptide-binding receptors and their paralogs. Then, all genes with no strong expression anywhere were removed (genes never expressed in more than 10% of cells of a specific cell type, across all cell types). Expression patterns of remaining GPCR genes were manually examined to generate lists of GPCRs expressed in non-sensory neurons.

### Identification of neuronal small secreted proteins (NSSPs)

In each species, we first selected all transcripts of genes for which the predicted translated coding sequence is of less than 200 amino acids. Sequences were filtered for the presence of a signal peptide using SignalP v.6.0 (Teufel *et al*. 2022) and the absence of a transmembrane domain using DeepTMHMM v.1.0 (Hallgren and al. 2022). All established neuropeptide precursor genes (and their paralogs) and all genes with domains detected through PFAM were removed except for DUF (“domain of unknown function”)-containing genes. Then, all cells in whole-animal datasets were classified into neuronal and non-neuronal cells, and normalized expression levels for all genes were pseudobulked according to this classification. Expression values were then examined in comparison to the set of neuropeptide precursor genes having a 1:1:1 in all three species. In each species, a gene from the filtered subset of small secreted genes was considered NSSP if its neuronal enrichment was ≥2 and if its normalized expression level in the nervous system was above the 10% decile value of the 1:1:1 neuropeptide genes (i.e., expression value above which 90% of the genomically conserved neuropeptide precursor genes are found). A slightly more stringent cutoff was used in *C. briggsae* (including 85% of the conserved neuropeptide precursor genes) as this made the cutoff more consistent relative to the observed expression distributions of the two other species. Resulting lists of genes are found in table S8.

### Statistics analyses and graphing

Neuropeptide network plots were generated using MATLAB (v24.1.0.2578822 (R2024a), The MathWorks Inc., Natick, MA). All other statistical analyses and plots were done in R V4.2.2. Statistical details for each analysis appear in figure legends. We mostly used the non-parametric Kruskal-Wallis test with Dunn’s correction for multiple comparisons (function dunn.test with method “bh”) and differences were considered statistically significant if p < 0.025 (α/2). When only two groups were compared, we used the Wilcoxon rank sum test (wilcox.test). For categorically-scored expression of fluorescent reporters out of all tested animals, we used Fisher’s test (fisher.test) with Bonferroni correction to multiple comparisons.

## Data availability

The raw sequencing data has been deposited in NCBI SRA (BioProject PRJNA851520). Annotated cell datasets (monocle3 objects) were deposited on Zenodo **10.5281/zenodo.14194526**. Analysis code is available at **10.5281/zenodo.14205686** (in progress).

## Acknowledgements

We thank Maria Antonietta Tosches, Itamar Lev, Stephanie Eder, Liesbet Temmerman, Giulio Valperga, Seth Taylor, and the colleagues in our labs for discussions about the project and feedback on the manuscript. We thank Qi Chen for her expertise in generating nematode strains in the lab. Other strains were provided by Christian Braendle, Erik Andersen and by the CGC (funded by the NIH, Office of Research Infrastructure Programs P40 OD010440). All research from the Department of Psychiatry at the University of Cambridge is made possible by the NIHR Cambridge Biomedical Research Centre and the NIHR East of England Applied Research Centre. The views expressed are those of the author(s) and not necessarily those of the NHS, the NIHR or the Department of Health.

## Funding

This work was funded by the Howard Hughes Medical Institute and the NIH grants RO1 NS039996 & NIH RO1 NS100547 (to OH); the NIH grant K99 HG010369 and Israel Science Foundation grant 2023/20 (to EBD); the Medical Research Council grant MC-A023-5PB91 (to WRS); the MQ Transforming Mental Health grant MGF17_24 (to PEV); and a postdoctoral fellowship from the Evelyn Gruss Lipper charitable foundation (to IAT).

## Author contributions

IAT, EBD and OH designed research. EBD generated scRNA-seq libraries. IAT, EBD, LTG analyzed data. IAT, LTG, KSS generated strains and performed experiments. LRS generated the neuropeptidergic connectomes. IB, PEV, WRS, EBD, OH supervised research. IAT and OH wrote the manuscript with input from all authors.

## Competing interests

EBD is an employee of Illumina Inc. All other authors declare they have no competing interests.

## Notes

### Competing Interest Statement

Eyal Ben-David is an employee of Illumina Inc. All other authors declare they have no competing interests.

