## Supplemental text and Figures for "Molecular patterns of evolutionary changes throughout the whole nervous system of multiple nematode species"

### Supplementary Text

<u>Detected differences in neurotransmitter-identity genes</u>

***eat-4/VGLUT* in DVA neurons**.

The tail neuron DVA, which is a dominantly-cholinergic neuron, was recently found to also express the vesicular glutamate transporter *eat-4* at low expression levels in *C. elegans* (WANG *et al.* 2024). We also detect dim but consistent *eat-4* expression in *C. elegans* DVA, while such expression is never detected in DVA neurons of *C. briggsae* or *C. tropicalis*.

***eat-4/VGLUT* in PVQ neurons**.

Our CRISPR reporter alleles highlighted expression of *eat-4* in PVQ neurons of *C. briggsae* and *C. tropicalis*. In *C. elegans*, *eat-4* expression in PVQ is detected in scRNA-seq datasets, in fosmid-based expression constructs and promoter-based constructs (SERRANO-SAIZ *et al.* 2013; TAYLOR *et al.* 2021), but is undetectable when examining the CRISPR reporter allele.

***cat-1* in RIR, CAN and VC4-5 neurons.**

The *cat-1/VMAT* reporter allele is consistently expressed in RIR, CAN and the VC4-5 neurons in *C. elegans*, but is OFF in these neurons in *C. briggsae* and *C. tropicalis*. In all three species, biosynthetic enzymes or uptake transporters for the known conventional monoaminergic transmitters are absent in RIR, CAN and VC4-VC5.

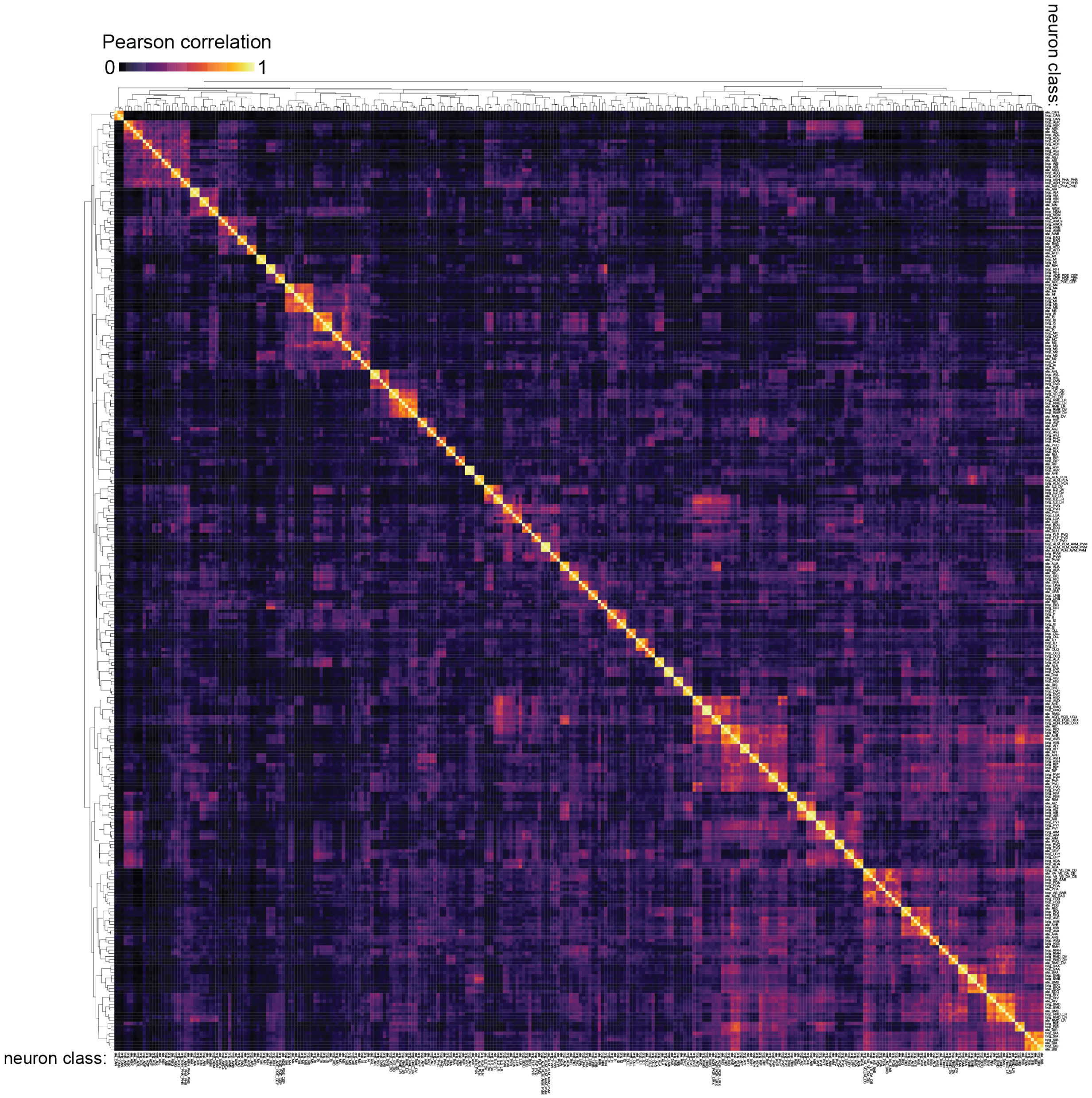


Fig. S1.
**Transcriptomic correlations of neuronal classes in three nematode species**. Hierarchical clustering and correlation matrix of pseudobulked normalized expression of 1,380 differentially-expressed 1:1:1 ortholog genes (top_markers score > 0.1 in at least one species and one neuron class). The triplets of homologous neuron classes clustered together across species. Correlation values are the same as those shown in Figure 1E, but neuron classes (x- & y-axes) are displayed according to the hierarchical clustering (not ordered by species & neuron class).

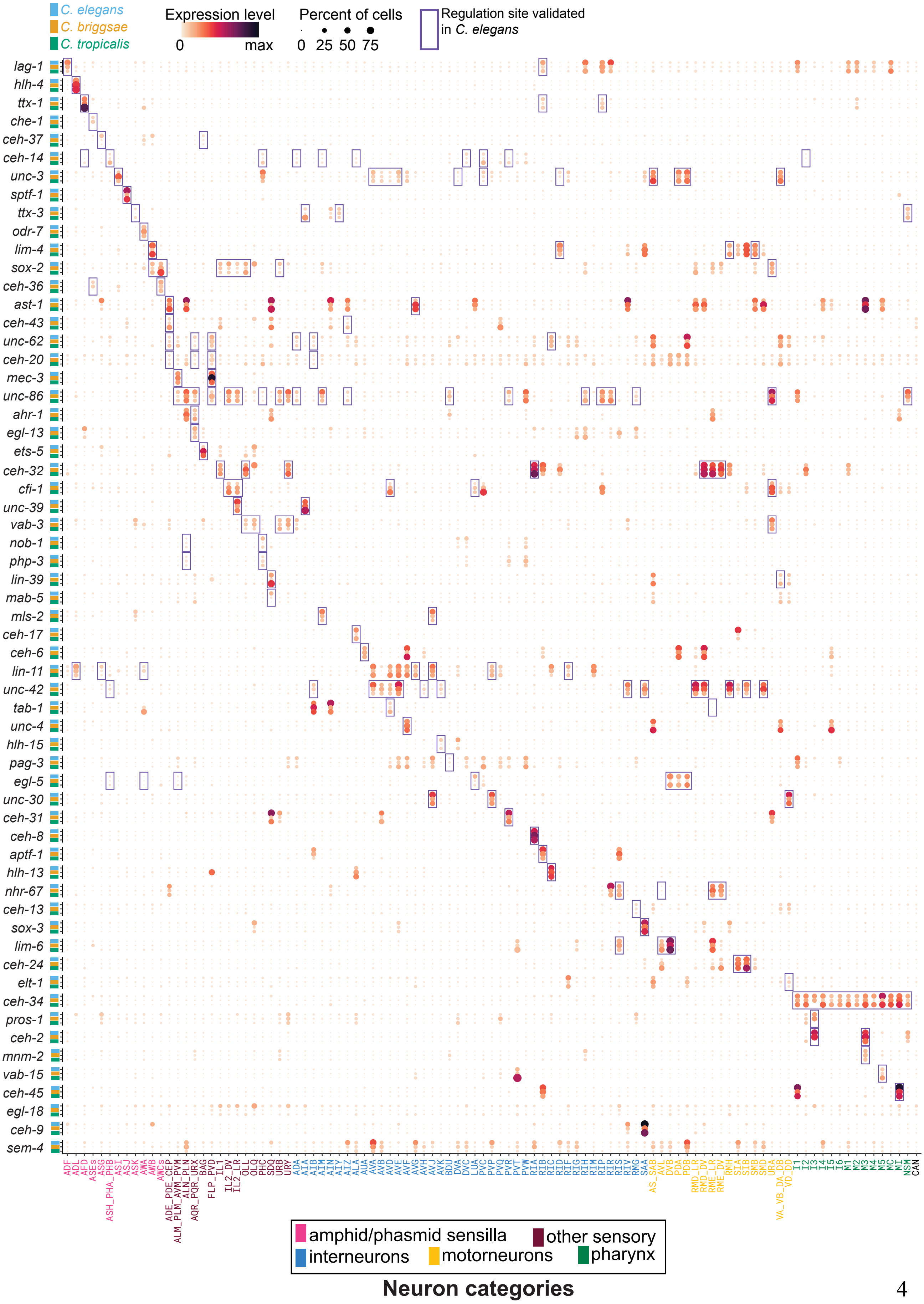

C. elegans
C. briggsae
C. tropicalis
Expression level
0
max
Percent of cells
0 25 50 75
Regulation site validated in C. elegans
lag-1
hlh-4
ttx-1
che-1
ceh-37
ceh-14
unc-3
sptf-1
ttx-3
odr-7
lim-4
sox-2
ceh-36
ast-1
ceh-43
unc-62
ceh-20
mec-3
unc-86
ahr-1
egl-13
ets-5
ceh-32
cfi-1
unc-39
vab-3
nob-1
php-3
lin-39
mab-5
mls-2
ceh-17
ceh-6
lin-11
unc-42
tab-1
unc-4
hlh-15
pag-3
egl-5
unc-30
ceh-31
ceh-8
aptf-1
hlh-13
nhr-67
ceh-13
sox-3
lim-6
ceh-24
elt-1
ceh-34
pros-1
ceh-2
mnm-2
vab-15
ceh-45
egl-18
ceh-9
sem-4
ADF
ADL
AFD
ASEs
ASG
ASH_PHA_PHB
ASI
ASJ
ASK
AWA
AWB
AWCs
ADE_PDE_CEP
ALM_PLM_AVM_PVM
ALN_PLN
AQR_PQR_URX
BAG
FLP_PVD
IL1
IL2_DV
IL2_LR
OLL
OLQ
PHC
SDQ
URB
URY
ADA
AIA
AIB
AIM
AIN
AIY
AIZ
ALA
AUA
AVA
AVB
AVD
AVE
AVF
AVG
AVH
AVJ
AVK
BDU
DVA
DVC
LUA
PVC
PVP
PVQ
PVR
PVT
PVW
RIA
RIB
RIC
RID
RIF
RIG
RIH
RIM
RIP
RIR
RIS
RIV
RMG
SAA
AS_SAB
AVL
DVB
PDA
PDB
RMD_LR
RMD_DV
RME_LR
RME_DV
RMH
SIA
SIB
SMB
SMD
URA
VA_VB_DA_DB
VD_DD
I1
I2
I3
I4
I5
I6
M1
M2
M3
M4
M5
MC
MI
NSM
CAN
amphid/phasmid sensilla
other sensory
interneurons
motorneurons
pharynx
Neuron categories

**Fig. S2.**

**Expression conservation of transcription factors composing the regulatory codes of neuronal identity in *C. elegans*.**

Cross-species expression dotplot of all 1:1:1 transcription factor orthologs functionally validated as identity specifiers in *C. elegans*. Nematode species (y-axis) and neuron class (x-axis) are color-coded according to legend. Dot size represents the fraction of cells expressing the gene in a given neuron class, color represents scaled average expression levels. Experimentally-validated sites of identity regulation by a given transcription factor in *C. elegans* are marked with a rectangle. A subset of genes and cells also appear in Figure 1F. Data for ASE and I3 neurons is available for *C. elegans* and *C. briggsae* datasets only. Data for *ceh-9* (that regulate PVN neurons) and for *egl-18* & *sem-4* (that regulate HSN neurons) are shown, but their corresponding neurons are not part of the dataset. PVN are born during L2, and HSN neurons acquire many of their class-specific identity features in the L4 larval stage.

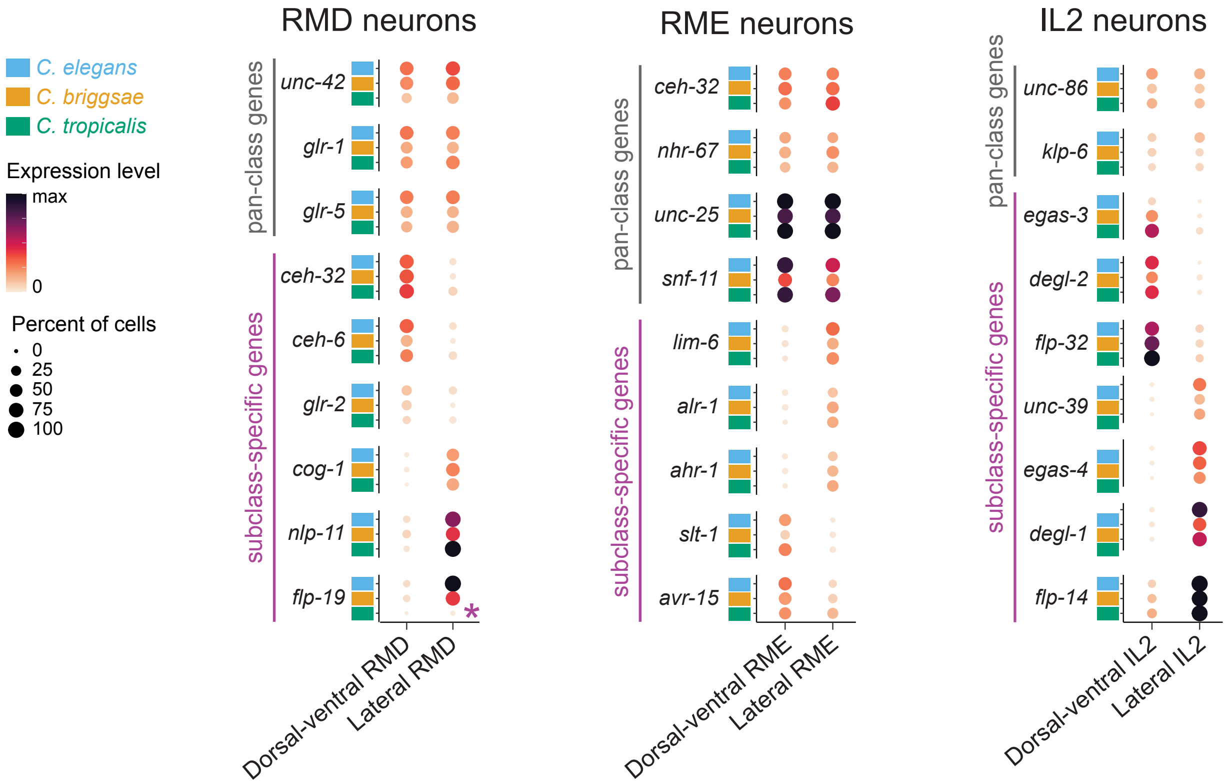


**Fig. S3.**
**Molecular distinctiveness of neuronal subclasses is conserved across nematode species**.
Cross-species expression dotplots of genes expressed in RMD-, RME- & IL2-class neurons. Represented subclasses are labeled in the x-axis. "Pan-class" genes (expressed in all subclasses) are marked in grey, and subclass-specific genes are marked in purple. Nematode species are color-coded according to legend. Dot size represents the fraction of cells expressing the gene in a given neuron class, color represents scaled average expression levels. Asterisk: expression of *flp-19*, a neuropeptide precursor gene expressed in RMD-lateral neurons, was absent specifically in *C. tropicalis.*

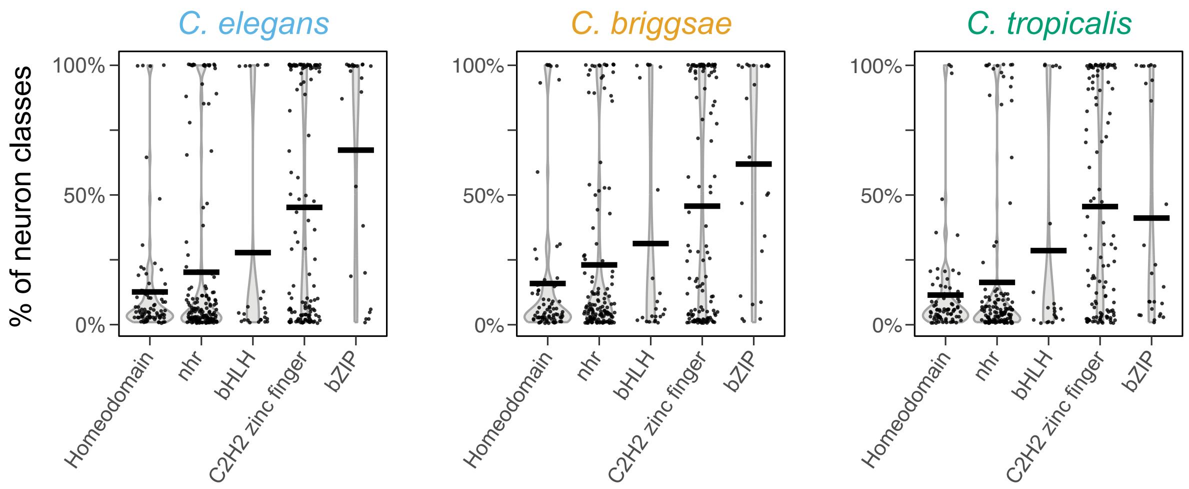


**Fig. S4.**
**Expression breadth of transcription factor families.**
Violin plots showing the proportion of neuron classes (y-axis) in which genes from different transcription factor families (x-axis) are expressed. Each dot represents one gene. Horizontal bars show average proportions per family. Homeodomain and NHR transcription factors tend to be expressed in fewer cell types in all three species. NHR genes, but not homeodomain genes, are heavily biased towards expression in sensory neurons (SURAL AND HOBERT 2021; TAYLOR *et al.* 2021).

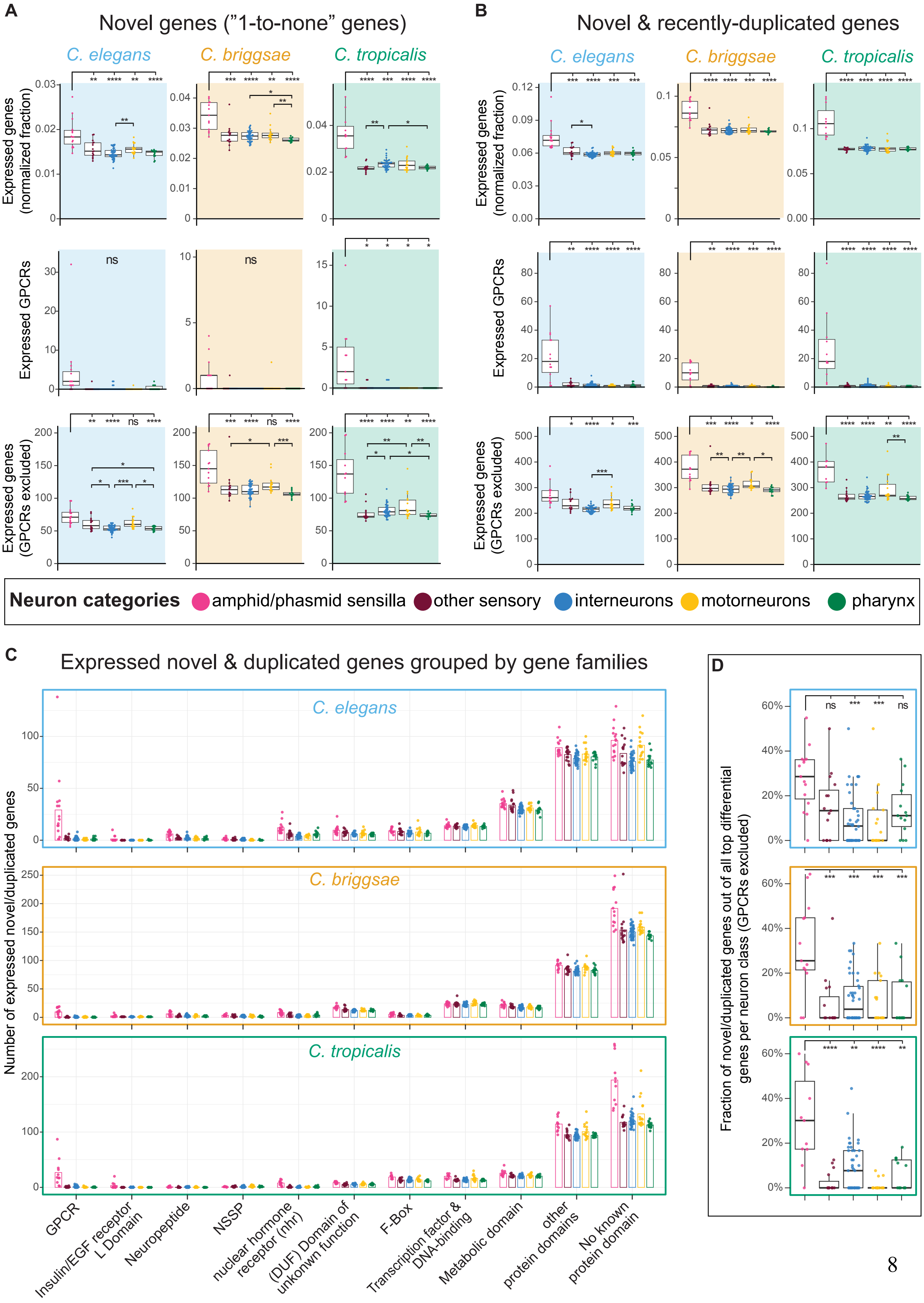

A
Novel genes ("1-to-none" genes)
B
Novel & recently-duplicated genes
C. elegans
C. briggsae
C. tropicalis
Expressed genes (normalized fraction)
Expressed GPCRs
Expressed genes (GPCRs excluded)
ns
Neuron categories
amphid/phasmid sensilla
other sensory
interneurons
motorneurons
pharynx
C
Expressed novel & duplicated genes grouped by gene families
Number of expressed novel/duplicated genes
GPCR
Insulin/EGF receptor L Domain
Neuropeptide
NSSP
nuclear hormone receptor (nhr)
(DUF) Domain of unkonwn function
F-Box
Transcription factor & DNA-binding
Metabolic domain
other protein domains
No known protein domain
D
Fraction of novel/duplicated genes out of all top differential genes per neuron class (GPCRs excluded)

**Fig. S5.**
**Enrichment of novel and duplicated genes in sensory neurons of the amphid and phasmid sensilla.**
(**A)** Expression of novel genes ("1-to-none" orthologs, y-axis) in different neuron classes and species (thresholded expression). Top panels: fraction of expressed novel genes normalized to the total number of expressed genes per cell type. Middle panel: number of novel GPCR genes expressed per cell type. Bottom panels: number of expressed genes after exclusion of all GPCRs from analysis. Neuron classes were grouped into functional categories (x-axis and color coded), each dot represents a single neuron class. Boxplots are Tukey-style.
(**B**) Expression of novel and recently-duplicated genes ("1-to-none", "1-to-many", "many-to-many" orthologs) in different neuron classes and species. Panels similar to (A).
(**C**) Number of expressed novel and recently-duplicated genes (y-axis) belonging to different gene families (x-axis) based on sequence homology or the presence of key protein domains. Neuron classes were grouped into functional categories, each dot represents a single neuron class, bars depict mean values for all neurons in category. Genes with no domains detected in PFAM are in the category "no known protein domain".
(**D**) Proportions of novel and duplicated genes out of all differentially-expressed genes per neuron class, grouped by neuron functional category. All GPCRs were excluded from the analysis.
Statistical tests: Kruskal-Wallis test with Dunn's correction. ****$P < 0.0001$, ***$P < 0.001$, **$P < 0.01$, *$P < 0.025$.

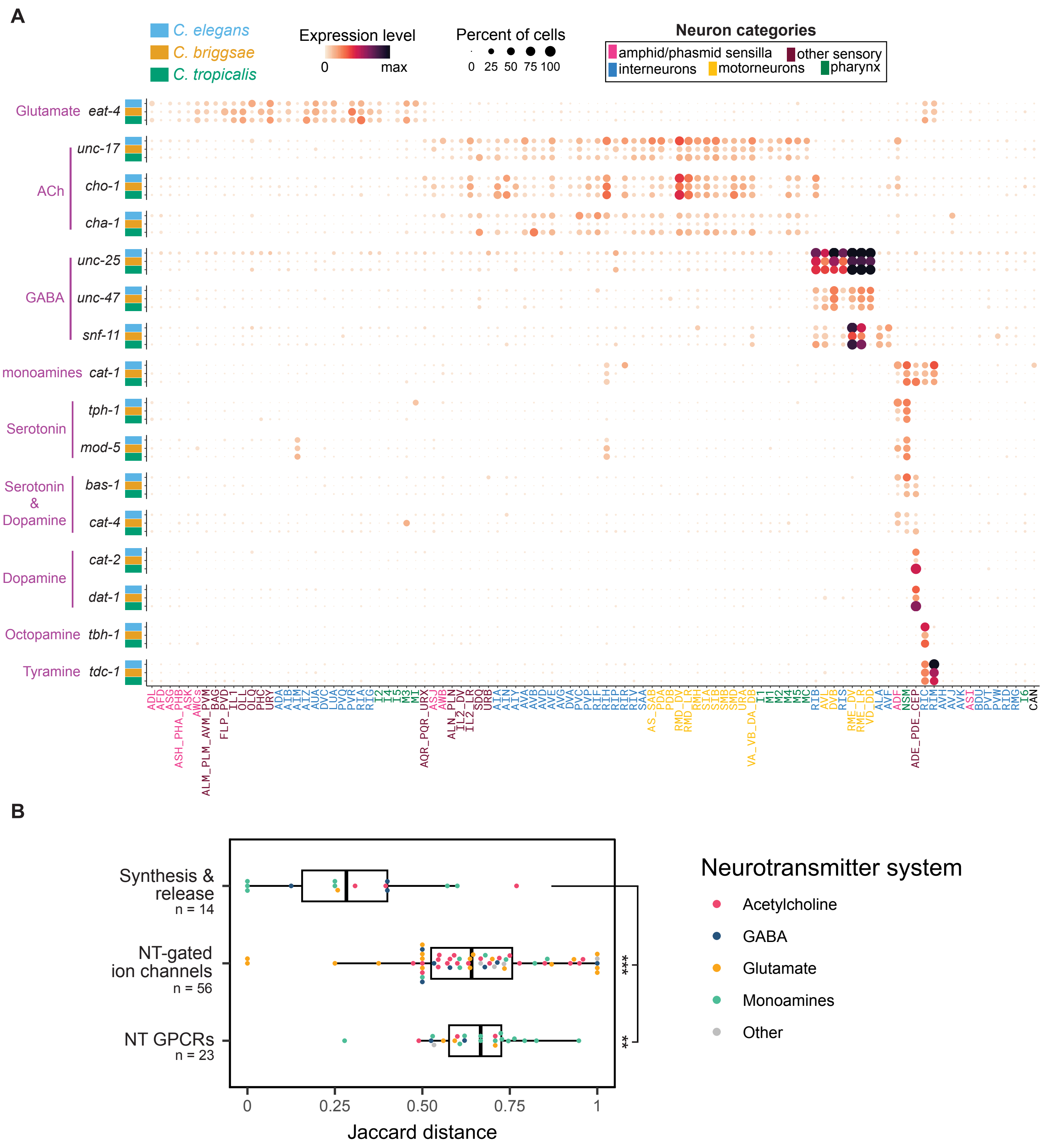


**Fig. S6.**

**Conserved expression of gene modules determining the identity of emitted neurotransmitters in homologous single neurons.**

(**A**) Cross-species expression dotplot of neurotransmitter synthesis genes, neurotransmitter vesicular transporter genes and neurotransmitter uptake transporter genes determining neurotransmitter release identity of neuronal cell classes. Data for *eat-4*, *unc*-17, *unc-25* & *cat-1* appear in Figure 3B and are shown again here for ease of visualization. Associated neurotransmitters are labeled in purple (y-axis). Ach: Acetylcholine; GABA: Gamma-aminobutyric acid. Nematode species (y-axis) and neuron class (x-axis) are color-coded according to legend. Dot size represents the fraction of cells expressing the gene in a given neuron class, color represents scaled average expression levels.

(**B**) Jaccard distances of 1:1:1 ortholog genes determining neurotransmitter release identity and receptivity to neurotransmitters. Each dot represents a gene, boxplots are Tukey-style. Data is a subset of the data already appearing in Figure 3A, but each gene is color-coded here according to the neurotransmitter system to which it belongs. Kruskal-Wallis test with Dunn's correction. ****P* < 0.001, ***P* < 0.01.

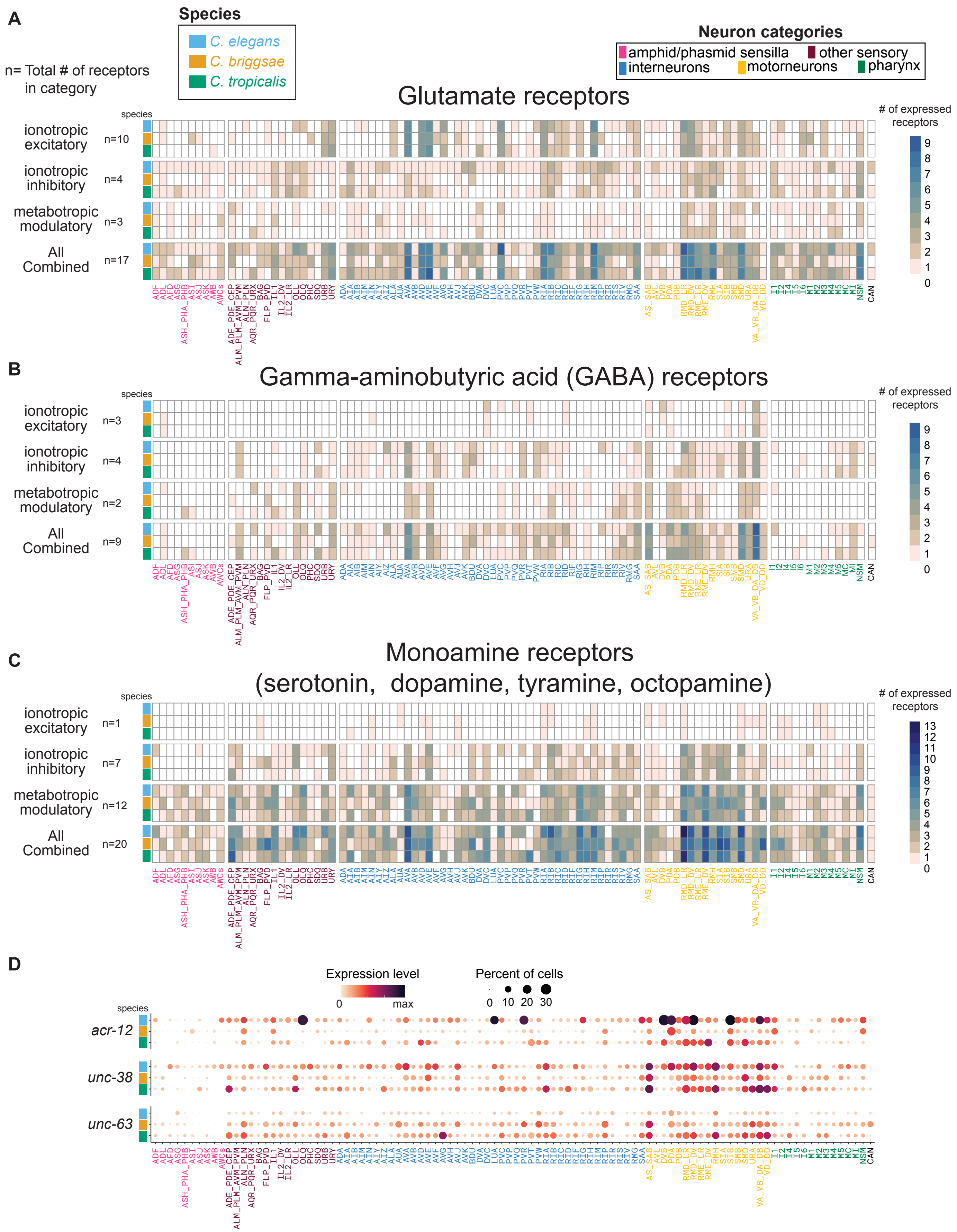
A
Species
C. elegans
C. briggsae
C. tropicalis
n= Total # of receptors in category
Neuron categories
amphid/phasmid sensilla
other sensory
interneurons
motorneurons
pharynx
Glutamate receptors
### of expressed receptors
ionotropic excitatory n=10
ionotropic inhibitory n=4
metabotropic modulatory n=3
All Combined n=17
B
Gamma-aminobutyric acid (GABA) receptors
ionotropic excitatory n=3
ionotropic inhibitory n=4
metabotropic modulatory n=2
All Combined n=9
C
Monoamine receptors (serotonin, dopamine, tyramine, octopamine)
ionotropic excitatory n=1
ionotropic inhibitory n=7
metabotropic modulatory n=12
All Combined n=20
D
Expression level
0
max
Percent of cells
0 10 20 30
acr-12
unc-38
unc-63

**Fig. S7.**
**Divergence in cell-type-specific expression of neurotransmitter receptors.**
(**A)** Heatmaps representing the number (color-coded) of glutamate receptors expressed in each neuron class (x-axis) across species (y-axis). Subtype of receptors (ionotropic excitatory, ionotropic inhibitory, metabotropic modulatory) are indicated to the left of heatmaps. N values indicate the total number of receptors included in the analysis for a given subtype.
(**B**) Heatmaps representing the expression of GABA receptors and 1:1:1 orthologs.
(**C**) Heatmaps representing the expression of monoamine receptors and 1:1:1 orthologs. Receptors for all known monoamines acting in *C. elegans* were considered combined together as a single category.
(**D**) Cross-species expression dotplot of the acetylcholine receptors *acr-12, unc-38* & *unc-63*. These receptors are broadly expressed throughout the nervous system. Nematode species (y-axis) and neuron class (x-axis) are color-coded according to legend. Dot size represents the fraction of cells expressing the gene in a given neuron class, color represents scaled average expression levels.

A

| NPP-GPCR | *C. elegans* | *C. briggsae* | *C. tropicalis* |
|---|---|---|---|
| NLP-13 DMSR-2 | pervasive | integrative | pervasive |
| FLP-1 DMSR-6 | local | local | broadcaster |
| FLP-4 DMSR-7 | broadcaster | pervasive | pervasive |
| FLP-7 DMSR-7 | pervasive | broadcaster | pervasive |
| FLP-12 DMSR-8 | local | broadcaster | broadcaster |
| FLP-4 EGL-6 | local | integrative | integrative |
| FLP-6 EGL-6 | local | integrative | integrative |
| FLP-8 FRPR-15 | local | local | broadcaster |
| FLP-3 FRPR-16 | broadcaster | local | local |
| FLP-1 FRPR-7 | local | broadcaster | broadcaster |
| FLP-4 FRPR-8 | local | integrative | integrative |
| FLP-21 NPR-1 | local | broadcaster | broadcaster |
| FLP-21 NPR-2 | broadcaster | local | local |
| FLP-15 NPR-3 | pervasive | integrative | pervasive |
| NLP-10 NPR-35 | local | integrative | integrative |
| NLP-13 NPR-41 | integrative | pervasive | no network |
| NLP-38 SPRR-2 | integrative | pervasive | pervasive |
| NLP-58 TKR-1 | local | broadcaster | integrative |
| NLP-58 TKR-2 | local | local | integrative |

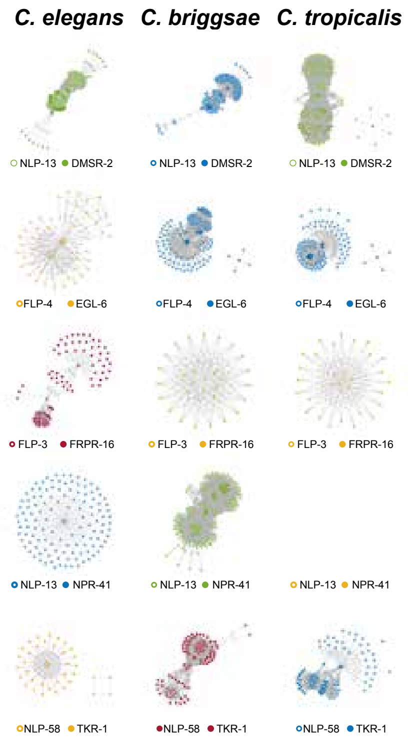


B

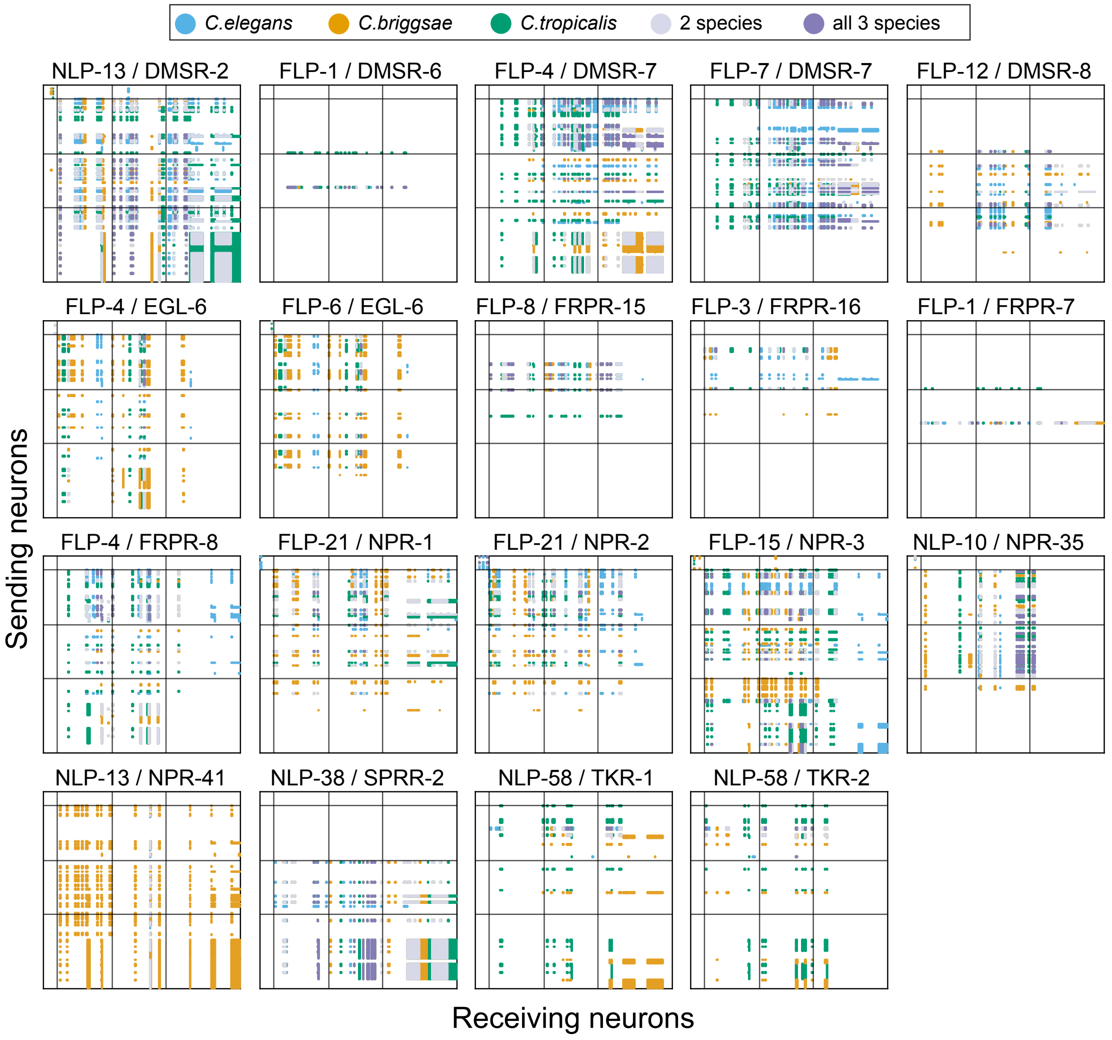

**Fig. S8.**
**Neuropeptide-receptor pairs displaying divergent network topologies across species.**
(**A**) 19 NPP-GPCR pairs displayed divergent topologies across species. Local networks (yellow) express the NPP and GPCR in ≤50 neurons. Pervasive networks (green) express both in ≥50 neurons. Broadcasting networks (red) express the NPP in ≤50 and the GPCR in ≥50. Integrative networks (blue) express the NPP ≥50 neurons and the GPCR in ≤50. Right: Graph visualizations of 4 NPP-GPCR networks across species.
(**B**) Adjacency matrix representation of the 19 individual NPP-GPCR pairs described above (short-range networks). Rows, columns and separations are similar to Figure 4E.

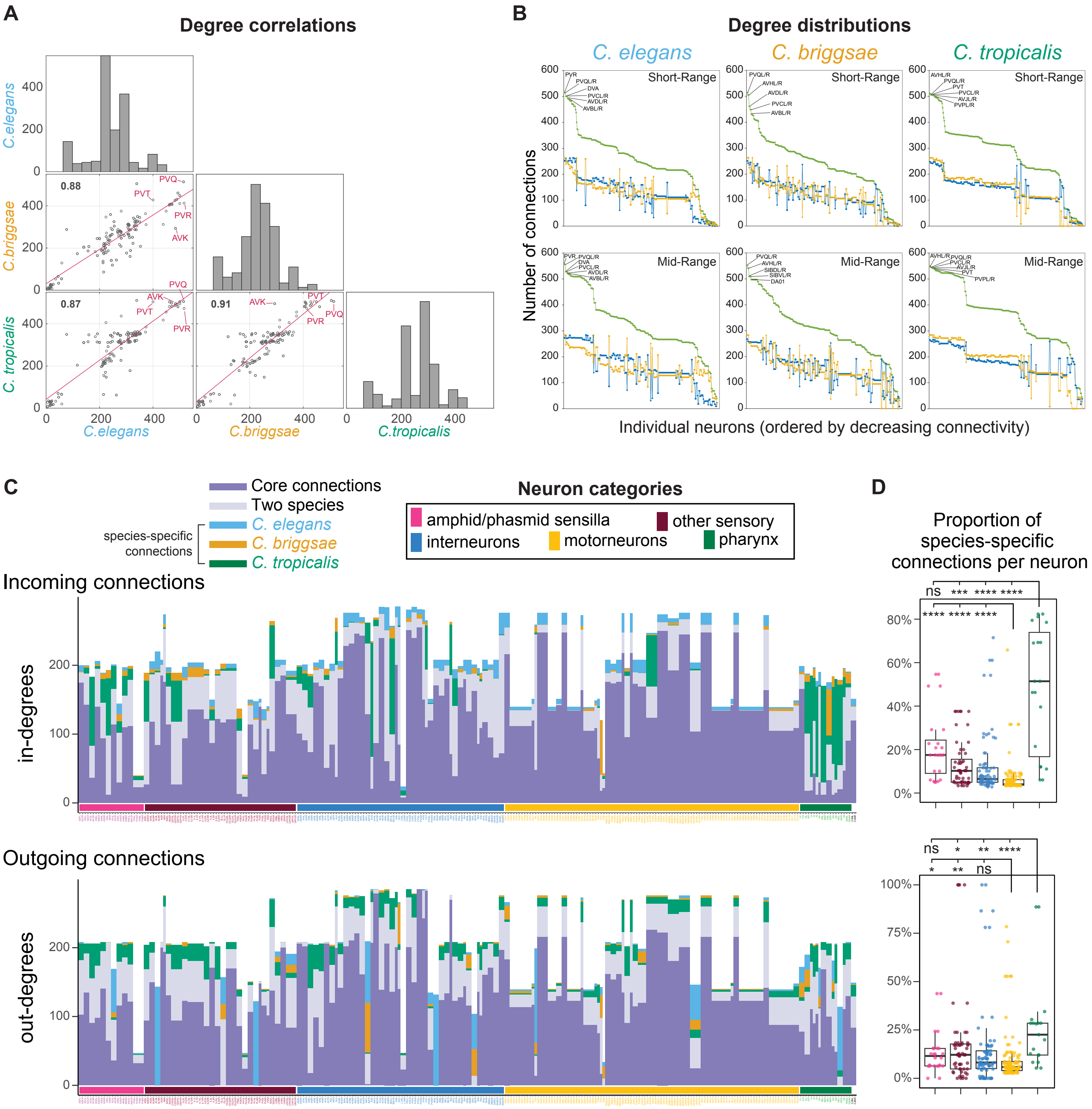


**Fig. S9.**

**Degree analysis in the neuropeptidergic networks.**

Peptidergic degree is defined as the number of incoming and outgoing connections per neuron (the sum of in-degrees and out-degrees).

(**A**) Pearson correlations of degrees between homologous neuron classes in pairs of species (short-range networks). The conserved peptidergic hubs are highlighted in red (AVK, PVR, PVQ, PVT).

(**B**) Distributions of degrees in the neuropeptidergic networks of three nematode species. Top panels: short-range networks. Bottom panels: mid-range networks. Degree (incoming plus outgoing connections) is shown in green, in-degree (incoming connections) in blue and out-degree (outgoing connections) in yellow. The 10 highest-degree hubs in each network are indicated.
(**C**) Total number of degrees (y-axis) in homologous neurons across species (x-axis). Top panel: in-degrees. Bottom panel: out-degrees. Bars are color-filled according to the subsets of core degrees and species-specific degrees of the neuron across species.
(**D**) Proportions of species-specific degrees (y-axis) per neuron classified by functional categories (colors and x-axis). Kruskal-Wallis test with Dunn's correction, ****$P < 0.0001$, ***$P < 0.001$, **$P < 0.01$, *$P < 0.025$.

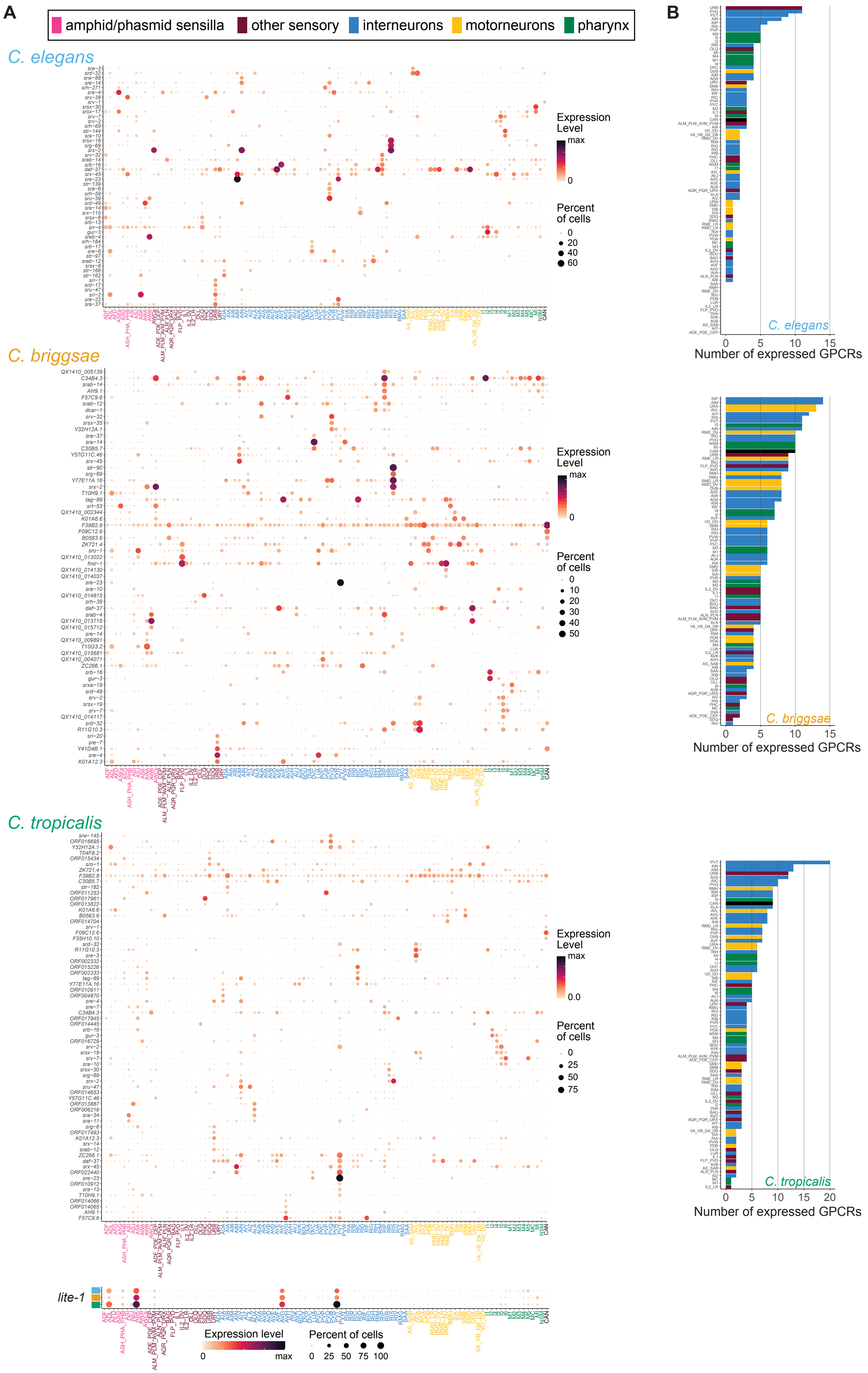
A
amphid/phasmid sensilla
other sensory
interneurons
motorneurons
pharynx
C. elegans
Expression Level
max
0
Percent of cells
C. briggsae
C. tropicalis
lite-1
Expression level
Percent of cells
B
Number of expressed GPCRs

**Fig. S10.**
**GPCRs expressed in non-sensory neurons.**
(**A**) Expression dotplots of GPCR genes that are expressed outside of the amphid/phasmid sensilla and have no sequence homology with neuropeptide- and neurotransmitter-binding GPCRs. 51 GPCRs pass the criteria in *C. elegans*, 63 in *C. briggsae* and 69 in *C. tropicalis*. In the *C. briggsae* and *C. tropicalis* panels, 1:1 orthologs of *C. elegans* genes appear with the *C. elegans* gene name (y-axis). Bottom: The light-responsive *lite-1* GPCR of the gustatory-receptor family is expressed at much higher levels than all other listed GPCRs, and is displayed separately. Dot size represents the fraction of cells expressing the gene in a given neuron class, color represents scaled average expression levels.
(**B**) Barplots indicating how many of these GPCRs are expressed in each neuron class. Bars are colored according functional categories, sensory neurons of the amphid and phasmid sensilla were excluded.

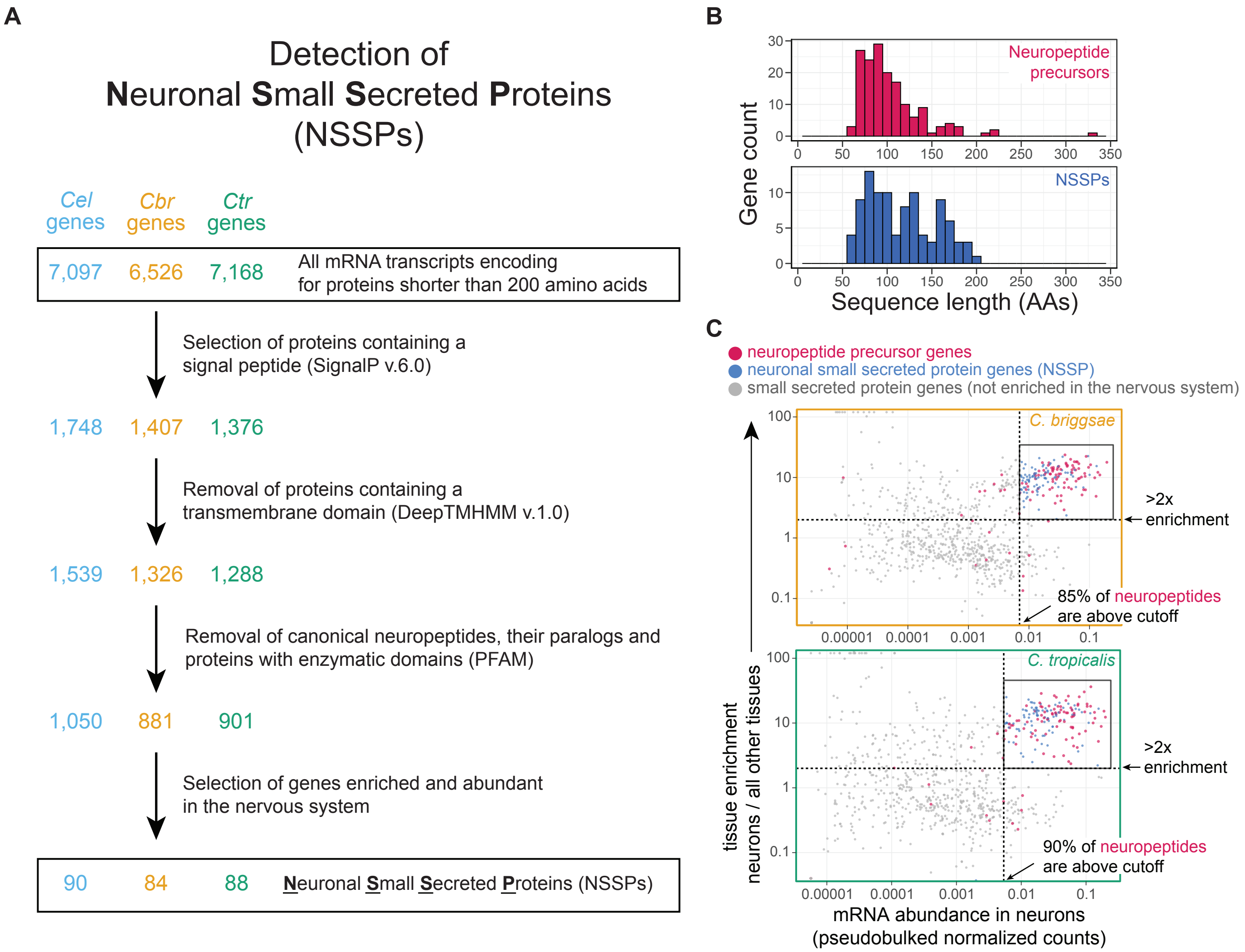


**Fig. S11.**
**Detection of Neuronal Small Secreted Proteins (NSSPs).**
(**A**) Diagram depicting the filtering process defining the pools of NSSPs. Numbers of genes remaining after each step are shown.
(**B**) Distributions of amino acid sequence lengths of all known *C. elegans* neuropeptide precursor genes (red) and newly-defined *C. elegans* NSSPs (blue). Longest mRNA isoform for each gene was used. Only 4/161 (2.5%) of all neuropeptide genes encode for proteins longer than 200 amino acids.
(**C**) Neuronal enrichment (y-axis) and neuronal expression levels (x-axis) of genes encoding small secreted proteins in *C. briggsae* (top panel) and *C. tropicalis* (bottom panel). Each dot is a gene, neuropeptides (red) and NSSPs (blue) are colored. Dashed lines: cutoff criteria used to delineate NSSPs.

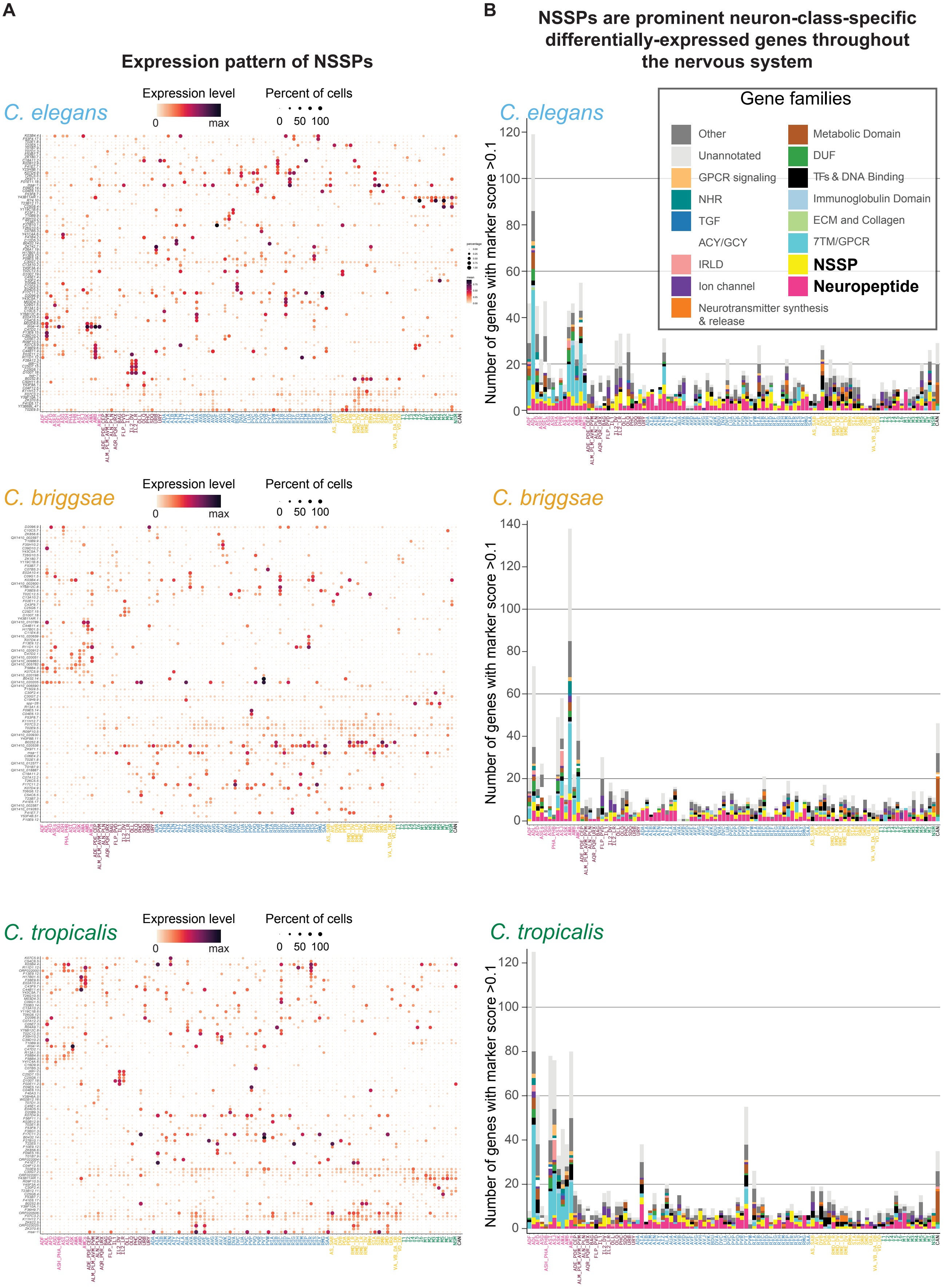
A
Expression pattern of NSSPs
C. elegans
Expression level
0
max
Percent of cells
0 50 100
C. briggsae
Expression level
0
max
Percent of cells
0 50 100
C. tropicalis
Expression level
0
max
Percent of cells
0 50 100
B
NSSPs are prominent neuron-class-specific differentially-expressed genes throughout the nervous system
Gene families
Other
Unannotated
GPCR signaling
NHR
TGF
ACY/GCY
IRLD
Ion channel
Neurotransmitter synthesis & release
Metabolic Domain
DUF
TFs & DNA Binding
Immunoglobulin Domain
ECM and Collagen
7TM/GPCR
NSSP
Neuropeptide
Number of genes with marker score >0.1

**Fig. S12.**
**NSSPs are a key distinctive molecular feature in most neuron classes.**
(**A**) Expression dotplots of NSSP genes in *C. elegans* (n=91 genes), *C. briggsae* (n=84) and *C. tropicalis* (n=88). In the *C. briggsae* and *C. tropicalis* panels, 1:1 orthologs of *C. elegans* genes appear with the *C. elegans* gene name (y-axis). Dot size represents the fraction of cells expressing the gene in a given neuron class, color represents scaled average expression levels.
(**B**) Barplots representing the numbers (y-axis) of differentially-expressed genes (*top_markers* score >0.1) expressed in each neuron class (x-axis). Genes with high marker scores tend to be abundantly and specifically expressed in one or few neuron classes. The fraction of genes belonging to specific gene families are colored according to legend. Neuropeptides (**magenta**) and NSSPs (**yellow**) appear among the most differentially-expressed genes in a majority of neuron classes throughout the nervous system. “NHR”: nuclear hormone receptor; “TGF”: Transforming Growth Factor family; “ACY/GCY” adenylate and guanylyl cyclases; “IRLD”: insulin/EGF receptor L Domain family; “DUF”: Domain of Unknown Function, “ECM”: extracellular matrix, “Unannotated”: genes with no domains detected by PFAM.

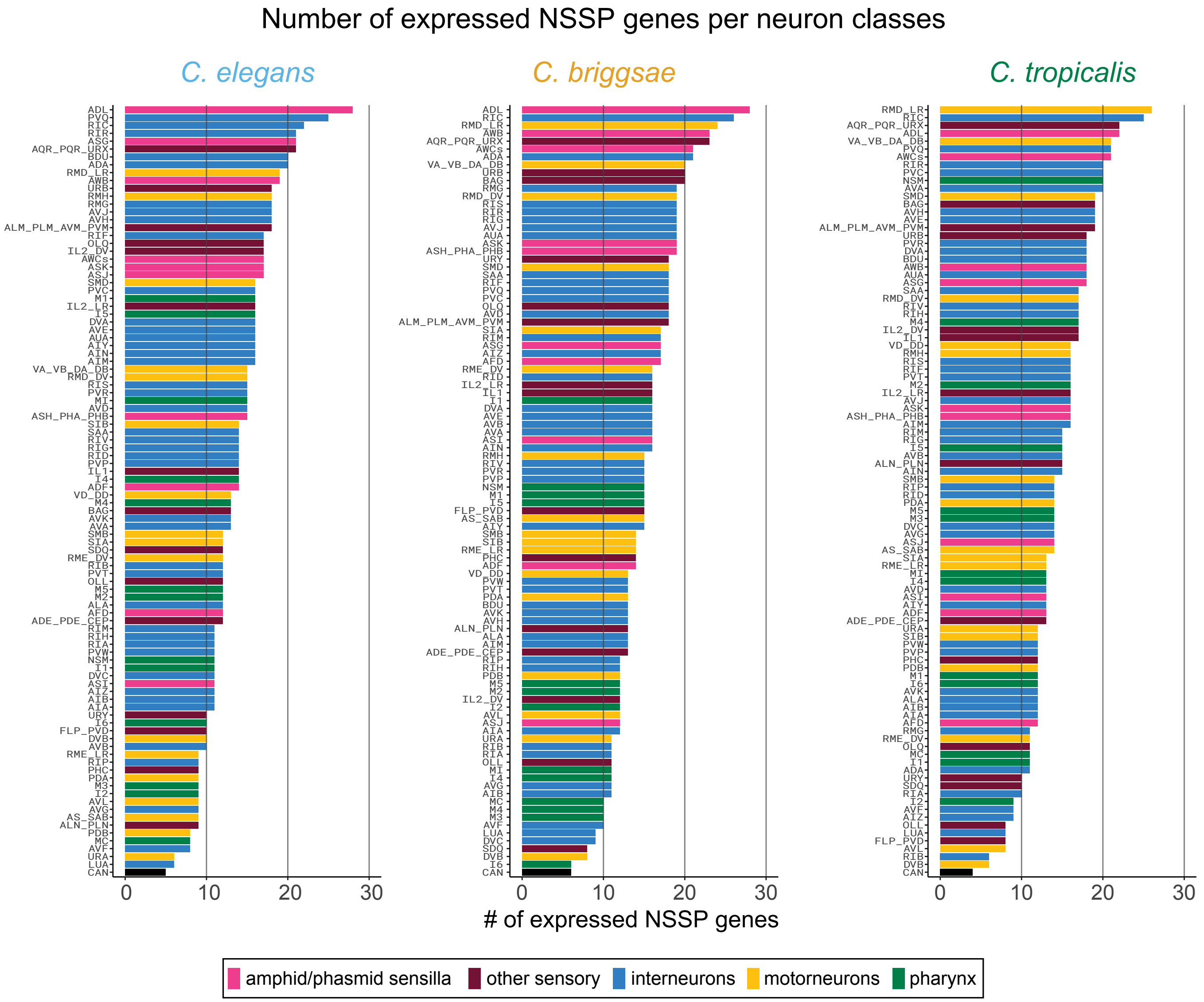


**Fig. S13.**
**Number of expressed NSSP genes in each neuron class.**
Barplots showing how many NSSPs are expressed in each neuron class. Bars are colored according functional categories of neurons. *Caenorhabditis* species indicated above plots.

### Supplementary Tables

**Table S1.**
**scRNA-seq summary statistics and cell annotations for three nematode species.**

**Table S2.**
**Top differentially-expressed genes in all neuronal and non-neuronal cell types.**
Output of the function top_markers from monocle3. Provides information about genes expressed in a cell-type-specific manner.

**Table S3.**
**Experimentally-validated "regulatory code" of *C. elegans* neuron classes.**
Table depicting all transcription factors regulating terminal identity markers in at a least one neuron class in *C. elegans*, based on experimental evidence.

**Table S4.**
**Ground truth data for expression thresholding and Jaccard distances for all 1:1:1 ortholog triplets**.

**Table S5**.
**List and information about all neurotransmitter receptors in *C. elegans* and receptors included in the analysis.**

**Table S6.**
**Wireless neuropeptidergic connectomes of three nematode species**.
Weighted networks and degree calculations for all neurons.

**Table S7.**
**GPCR genes and families**.
Lists of GPCR genes according to ligand family (neuropeptide-binding, neurotransmitter-binding, other). Filtered list of non-NP and non-NT GPCRs that are expressed in neurons that are not sensory neurons of the amphid and phasmid sensilla. Analysis of number of expressed "other" GPCR genes per neuron class in each species.

**Table S8**.
**Neuronal small secreted proteins (NSSPs)**.
Lists of all genes fulfilling the NSSP criteria in each species. Analysis of number of expressed NSSP genes per neuron class in each species.

**Table S9**.
**Information about strains, CRISPR reagents and plasmids generated and used in this study.**

**Table S10.**
**Orthology and protein family information for all genes.**

### Supplementary Materials References


Serrano-Saiz, E., Richard J. Poole, T. Felton, F. Zhang, Estanisla D. De La Cruz *et al.*, 2013 Modular Control of Glutamatergic Neuronal Identity in C. elegans by Distinct Homeodomain Proteins. Cell 155**:** 659-673.

Sural, S., and O. Hobert, 2021 Nematode nuclear receptors as integrators of sensory information. Curr Biol 31**:** 4361-4366 e4362.

Taylor, S. R., G. Santpere, A. Weinreb, A. Barrett, M. B. Reilly *et al.*, 2021 Molecular topography of an entire nervous system. Cell 184**:** 4329-4347 e4323.

Wang, C., B. Vidal, S. Sural, C. Loer, G. R. Aguilar *et al.*, 2024 A neurotransmitter atlas of C. elegans males and hermaphrodites. Elife 13.
